# Bone Response to Mechanical Loading is Highly Heritable and Correlates with Osteocyte Lacunar Morphology in Genetically Diverse Mouse Strains

**DOI:** 10.64898/2026.10.30.763214

**Authors:** Nicole Migotsky, Syeda N. Lamia, John T. Shuster, Surabhi Kumar, Charles R. Farber, Matthew J. Silva

## Abstract

Skeletal loading is a key regulator of bone formation. Studies using in vivo loading of rodent bones have established many mechanical and biological factors that regulate bone’s response to loading. However, most studies have been performed in a small number of inbred rodent strains, and there are few data on effects of genetic background on the loading response. In the current study we employed two murine models of genetic diversity, the Diversity Outbred (DO) and their eight founder inbred strains, to assess the influence of genetic background on skeletal response to tibial loading. Loading-induced periosteal bone formation rates and formed bone volumes varied significantly among inbred mouse strains. DO mice, which each carry a unique mixture of alleles of the inbred founder strains, were more responsive to loading, on average, than the the inbred founder strains. Additionally, bone’s response to mechanical loading was highly heritable; broad-sense heritability for measures of periosteal formation were near or above 80%. Furthermore, osteocyte lacunar morphology correlated with loading response across inbred strains. Inbred 129S1 mice had the most elongated lacunae and exhibited the highest magnitude of response on both periosteal and endocortical surfaces. 129S1 also demonstrated a robust transcriptional response to loading, although the pathways induced by loading were largely conserved with lab-standard C57Bl/6. Together, these findings demonstrate that genetic background significantly contributes to bone’s response to loading and establish DO mice as a platform for discovery of genes that regulate skeletal mechanoresponsiveness.

## Introduction

The regulation of bone (re)modeling by physical forces is a fundamental feature of skeletal physiology (1). Mechanical loading is a potent bone anabolic stimulus, and pre-clinical studies of this mechanobiological process can be used to identify genes and pathways that influence bone mass (2, 3). Skeletal responsiveness to loading is conserved across animal species (e.g., mice, rats, and humans (4–6)), although most of what we know about the biological mechanisms underlying this response comes from studies in C57Bl/6 (B6) mice or transgenic mice on a B6 background (7). More than 20 years ago, a significant influence of genetic background on bone mechanoresponsiveness was recognized (8, 9), and one study reported that the response to loading was highly heritable (H^2^=0.7-0.8) (10). However, these earlier studies were done using only two or three strains of closely related inbred mice, and this topic has not been revisited.

Analysis of more than 100 inbred mouse strains (*Mus musculus*) revealed that the genomes of classical inbred laboratory mouse strains (e.g., B6) are derived primarily from the subspecies *M. m. domesticus* and have limited genetic diversity (11). The inclusion of wild-derived strains from other subspecies (including *M. m. musculus* and *M. m. castaneus*) can add substantially to the diversity of a population. Researchers have identified eight inbred strains that collectively capture 89% of the variation in the mouse genome (12): five classical strains (A/J, C57BL/6J, 129S1/SvImJ, NOD/LtJ, NZO/HlLtJ) and three wild-derived strains (CAST/EiJ, PWK/PhJ, WSB/EiJ). We recently reported large inter-strain variation and high heritability (>80%) in a majority of cortical bone traits measured across these eight inbred strains, including cortical area and ultimate bending force (13). Moreover, Friedman et al. used these strains and showed that the magnitude of bone loss due to limb unloading is heritable (14, 15). Taken together, these studies support the use of these eight inbred strains to robustly determine the heritability of bone’s response to increased mechanical loading.

These same eight inbred strains were used as the founders for creation of an outbred population wherein each mouse is genetically unique, the Diversity Outbred (DO) mouse resource (16). DO mice can better model the genetic heterogeneity that is seen in humans, and have been used to discover novel genes regulating bone mass and other quantitative traits (17, 18). Thus, they represent a potential tool for discovery of new genes that regulate bone adaptation to mechanical loading. However, the prerequisites for such an endeavor are lacking, i.e., evidence that bone’s response to loading varies across the founder strains, and that the DO mice exhibit a continuous range of loading responses.

Sexual dimorphism in skeletal biology and disease is recognized as an important, but understudied research area (19, 20). Both female and male B6 mice exhibit increased bone formation in response to increasing mechanical strain (21). According to a recent review, more than 70% of studies of bone loading in wildtype mice used females (mostly B6), and only 5% included both sexes (7). In studies of knockout mice, inclusion of males and females is more common (∼20%) (7), but sex is rarely treated as an independent experimental factor. Even when statistical comparisons of females and males are reported, the influence of sex can depend on genotype and skeletal site (22). Overall, there is a need for more data to better understand whether, or in what context, the response to bone loading depends on sex.

Osteocytes are highly mechanoresponsive and as such are likely to contribute to the regulation of bone’s response to mechanical loading (1, 23). Fluid flow through the osteocyte lacunar-canalicular network (LCN) is considered the mechanism whereby external bone loading is converted to a biophysical stimulus at the osteocyte level (24, 25). Recent computational studies predict that lacunar shape affects fluid velocities in the LCN and the induced cellular strains, with more elongated lacunae producing higher local velocities and cell strains (26, 27). Interestingly, we recently reported that morphological traits of osteocyte lacunae vary across the eight inbred strains, with lacunar number and aspect ratio (a measure of elongation) having high heritabilities (H^2^ = 0.76-0.97) (13). This leads us to hypothesize that differences in lacunar morphology may correlate with differences in responsiveness to loading between mouse strains.

Our objective study was to investigate the effect of genetic diversity on bone response to mechanical loading. We applied axial tibial loading to females and males from two genetically diverse cohorts of mice: the eight inbred founder strains, and DO mice. Responses to loading were assessed using serial microCT, dynamic histomorphometry, and bulk RNAseq of bone tissue (Fig. 1). We asked: 1) Does loading-induced bone formation vary between the eight inbred founder mouse strains? 2) Does loading-induced bone formation vary between females and males of these strains? 3) How does the loading response of mice from the Diversity Outbred (DO) population compare to the response of the eight founder strains? 4) Does loading-induced bone formation correlate with homeostatic bone traits in inbred mice, including osteocyte lacunar morphology?

**Fig. 1:**
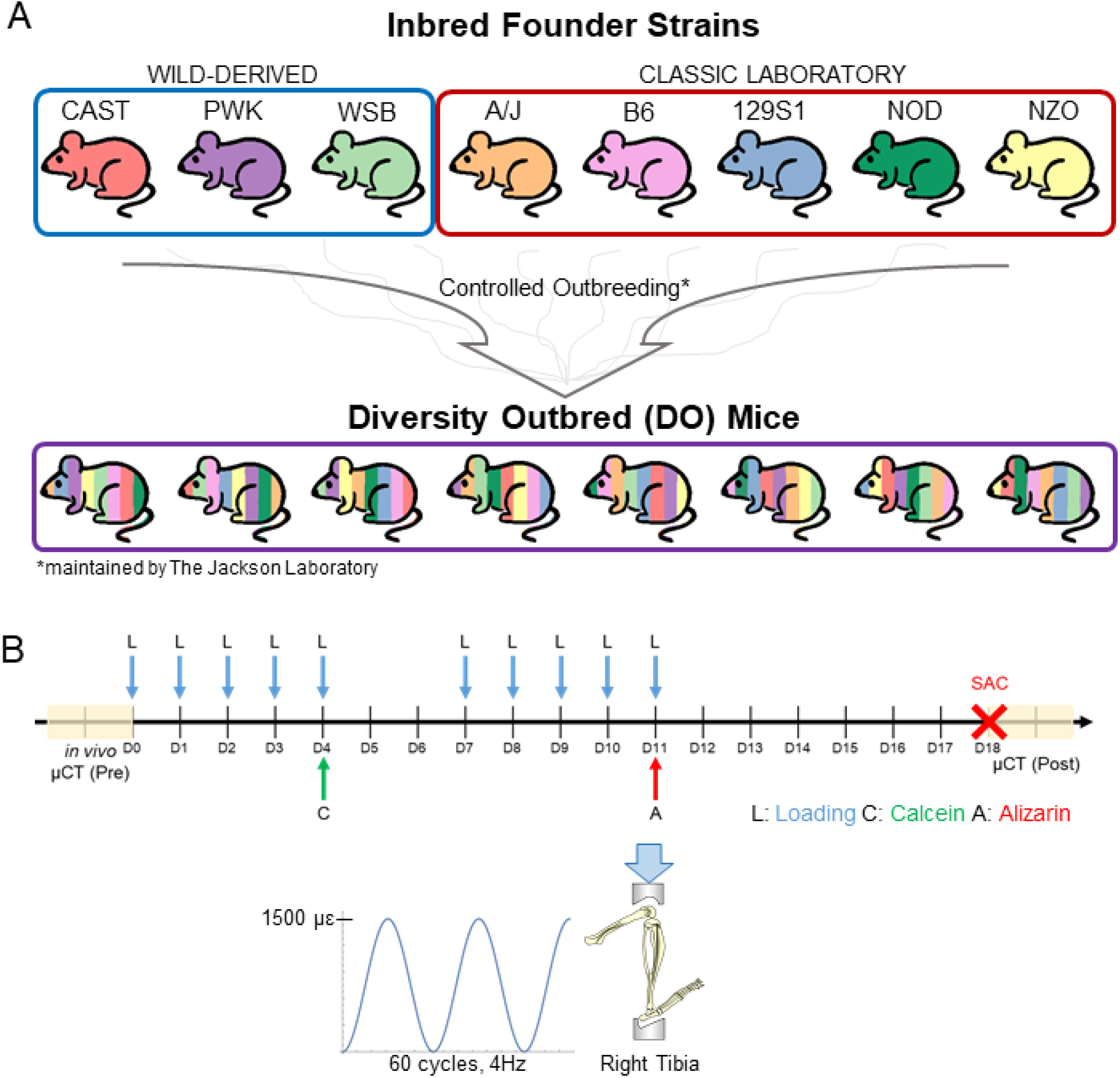
Overview of experimental groups and timeline. (A) Two mouse models of genetic diversity were used, eight inbred mouse strains and an outbred stock. The inbred strains were the founders used by The Jackson Laboratory to create the diversity outbred (DO) population. After controlled outbreeding of the founder lines, each DO mouse is genetically unique. Loading was performed on 5-month-old female and male mice from the eight Inbred Founder strains and the DO mice. (B) The right tibiae of mice were loaded 5 days per week for 2 weeks. At the end of each week of loading mice were injected with either calcein or alizarin. µCT scans were taken before and after loading to track change in morphology.

## Results

### Loading-Induced Changes in Bone Morphology Vary Significantly with Mouse Strain

Adult mice from Inbred Founder and DO populations were subjected to daily tibial loading for 2 weeks to stimulate cortical bone formation (Fig. 1B). Loading forces were adjusted to engender equivalent local bone strain magnitude at the mid-diaphysis in each mouse (Suppl. Fig. S1). To assess the effect of loading, we calculated the loading-induced changes in cortical bone morphometry outcomes using serial microCT (Fig. 2A). Changes in bone area due to loading varied significantly with mouse strain in a sex-dependent manner (strain: p<0.0001; strain-sex interaction: p<0.0001; Fig. 2B). Ten of the 16 strain/sex groups had a positive, significant response to loading (i.e., loading-induced change different from zero). Inbred strain 129S1 had the largest increase in bone area for both sexes (F: 21% increase in loaded limb, M: 20%; Suppl. Tables S8,S9). In comparison, B6 mice had moderate increases (F: 8.4%, M: 11.7%), while the morphology of WSB and A/J mice did not change significantly with loading. The main effect of sex was not significant, although significant sex differences were seen in CAST and PWK mice: CAST females were less responsive than CAST males, but PWK females were more responsive than PWK males. Similar changes were observed in other measures of cortical bone size (Tt.Ar, Ct.Th, pMOI; Suppl. Fig. S4). A significant loading-induced change in medullary area was observed in only one of 16 groups (NZO females), indicating a marginal effect of loading on the endocortical surface (Fig. 2D).

**Fig. 2:**
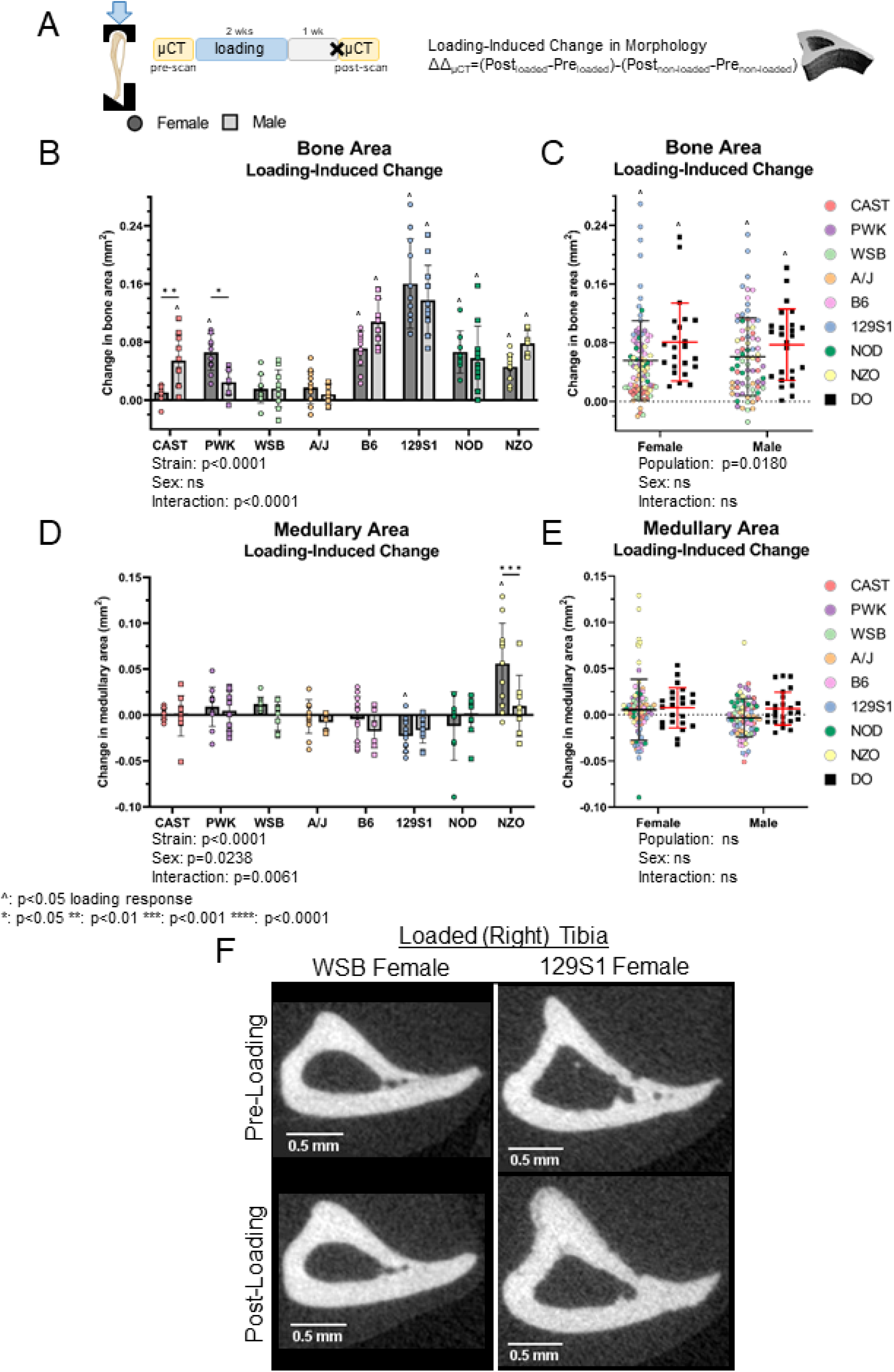
Loading-induced changes in bone morphology vary with mouse strain. (A) Tibiae were unilaterally loaded daily for 2 wks, and both nonloaded and loaded tibiae were microCT-scanned at the beginning (baseline) and end of the study period. Relative changes with time were calculated for each limb with respect to baseline (Δ), and then relative changes due to loading were calculated as differences between limbs (ΔΔ). (B) Relative changes in bone area of eight inbred strains and (C) inbred strains (pooled) and diversity outbred (DO) mice. (D) Relative changes in medullary area of inbred strains and (E) inbred strains (pooled) and DO mice. (F) Representative microCT images of WSB and 129S1 pre-and post-loading. Two-way ANOVA (sex and strain or population). Data shown as mean±SD.

Compared to the Inbred Founders, the DO population on average had a higher increase in bone area, total area and moment of inertia, indicating a greater response to loading (Fig. 2C, Suppl. Fig. S4A,C,E). The majority of values for DO mice fell within the range of the values of the Inbred Founders. There was no effect of sex on increases in bone parameters in either the pooled Inbred Founder population or the DO population. There were no changes in TMD due to loading in any Inbred Founder strain or the DO population (Suppl. Fig. S4G,H).

In summary, measures of bone size increased with loading in the majority of Inbred strains and in DO mice. Changes in morphology due to loading varied strongly between Inbred Founder strains, with only modest variations due to sex. The Diversity Outbred population exhibited a greater response to loading compared to the Inbred Founders.

### Loading-Induced Increases in Formed Bone Volume and Decreases in Resorbed Bone Volume Vary Significantly with Mouse Strain

To assess changes in bone formation and resorption on the periosteal and endocortical surfaces, we compared the μCT post-scan to the pre-scan using time-lapse analysis of individual voxels (Fig. 3A, Suppl. Fig. S3A). In response to loading, 11 of the 16 strain/sex groups had significantly increased formed bone volume on the periosteal surface (Fig. 3B). The magnitude of response varied significantly between strains in a sex-dependent manner, with both PWK and 129S1 females exhibiting larger increases in formation than males. Similar to changes in bone morphology, 129S1 mice had the greatest increase in periosteal formed bone (129S1 F: 19%, M: 15%; Suppl. Tables S6,S7) while WSB mice had a minimal response (significant only in females) (Fig. 3B,F). In addition, 8 of the 16 strain/sex groups had significantly decreased resorbed bone volume on the periosteal surface while no strains had increased resorption (Fig. 3D). The magnitude of decrease in resorption (avg. approx. 1%) was substantially less than the increase in formation (avg. approx. 6%) but also varied between strains in a sex-dependent manner. Only CAST mice showed a significant sex difference, with males having a larger decrease in resorption. Compared to the Inbred Founders, DO mice on average had a larger increase in formation on the periosteal surface, with all DO values within the range of the Inbred mice (Fig. 3C). Resorbed periosteal volume did not differ between the two populations (Fig. 3E).

**Fig. 3:**
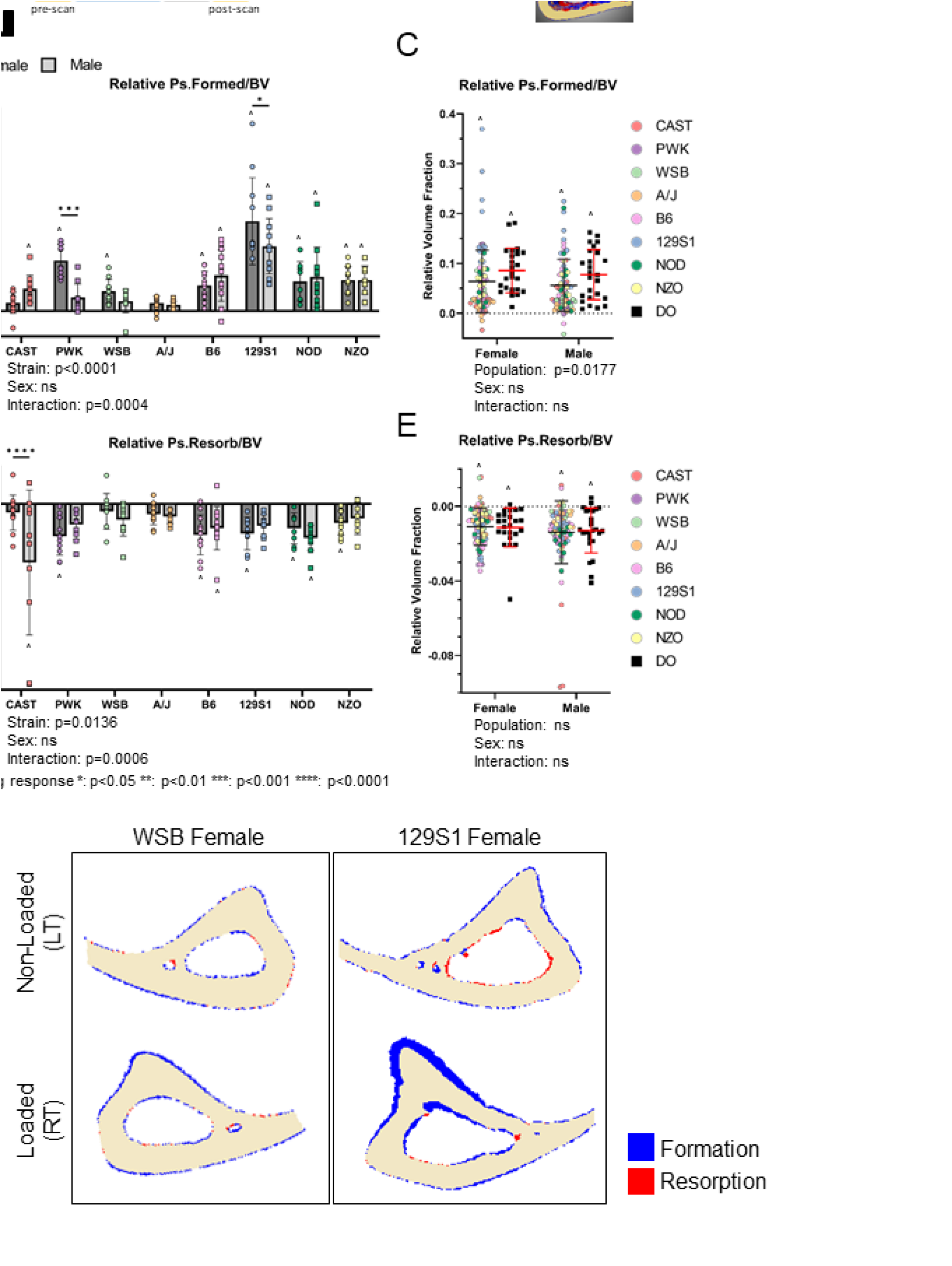
Loading-induced increases in periosteal formed bone volume vary with mouse strain and sex. (A) The pre- and post-loading tibia microCT scans were registered in 3D to obtain formed and resorbed bone volumes. Relative changes were calculated with respect to nonloaded tibia. (B) Relative periosteal (Ps.) formed volume fraction in eight inbred strains and (C) in inbred strains (pooled) and diversity outbred (DO) mice. (D) Relative periosteal resorbed volume fraction in inbred strains and (E) in inbred strains (pooled) and DO mice. (F) Representative registered images of WSB and 129S1 showing formed and resorbed voxels. Two-way ANOVA (sex and strain or population). Data shown as mean±SD.

On the endocortical surface, bone formation increased in only 4 of 16 strain/sex groups, and the magnitude of increase was less than on the periosteal surface (Suppl. Fig. S5A). Nonetheless, the variation between strains remained highly significant due to the increases in B6 and 129S1 strains (129S1 F: 5.1%, M: 3.8%). The effects of loading on endocortical resorbed bone were modest and reached significance in only two groups; NOD females had decreased bone resorption in response to loading while NZO females had increased resorption (Suppl. Fig. S5C). Comparing populations, no significant differences were observed between the Inbred Founders and DO mice for endocortical formed bone (Suppl. Fig. S5B), although DO mice had marginally higher increases in endocortical resorbed bone than Inbred mice (p=0.049).

In summary, time-lapse microCT analysis revealed significant strain-dependent variations in the amounts of formed and resorbed bone volume in response to loading, with only modest sex effects. 129S1 mice had the greatest amount of formed bone on both periosteal and endocortical surfaces. On average, the Diversity Outbred population had a greater periosteal response than the Inbred Founders.

### Increases in Bone Formation Rates Due to Loading Vary with Mouse Strain and Sex

We also assessed relative indices of loading-induced bone formation using classic 2D dynamic histomorphometry based on fluorochrome labels (Fig. 4A). On the periosteal surface, relative mineralizing surface (MS/BS), mineral apposition rate (MAR) and bone formation rate (BFR/BS) all varied significantly with mouse strain (Fig. 4B, Suppl. Fig. S6A,C). Ten or more of the 16 strain/sex groups had positive, significant responses to loading based on these periosteal indices. Mineralizing surface and mineral apposition rate, but not bone formation rate, had significant strain/sex interactions, although the only pairwise difference was in rPs.MS/BS for PWK (female > male). Inbred strain 129S1 had the largest increase in bone formation indices on the periosteal surface (Ps.BFR/BS; F and M: 11-fold increase, Suppl. Tables S4,S5). Compared to the Inbred Founders, the DO population on average had larger increases in periosteal bone formation rate, with all DO values within the range of the Inbred mice (Fig. 4C).

**Fig. 4:**
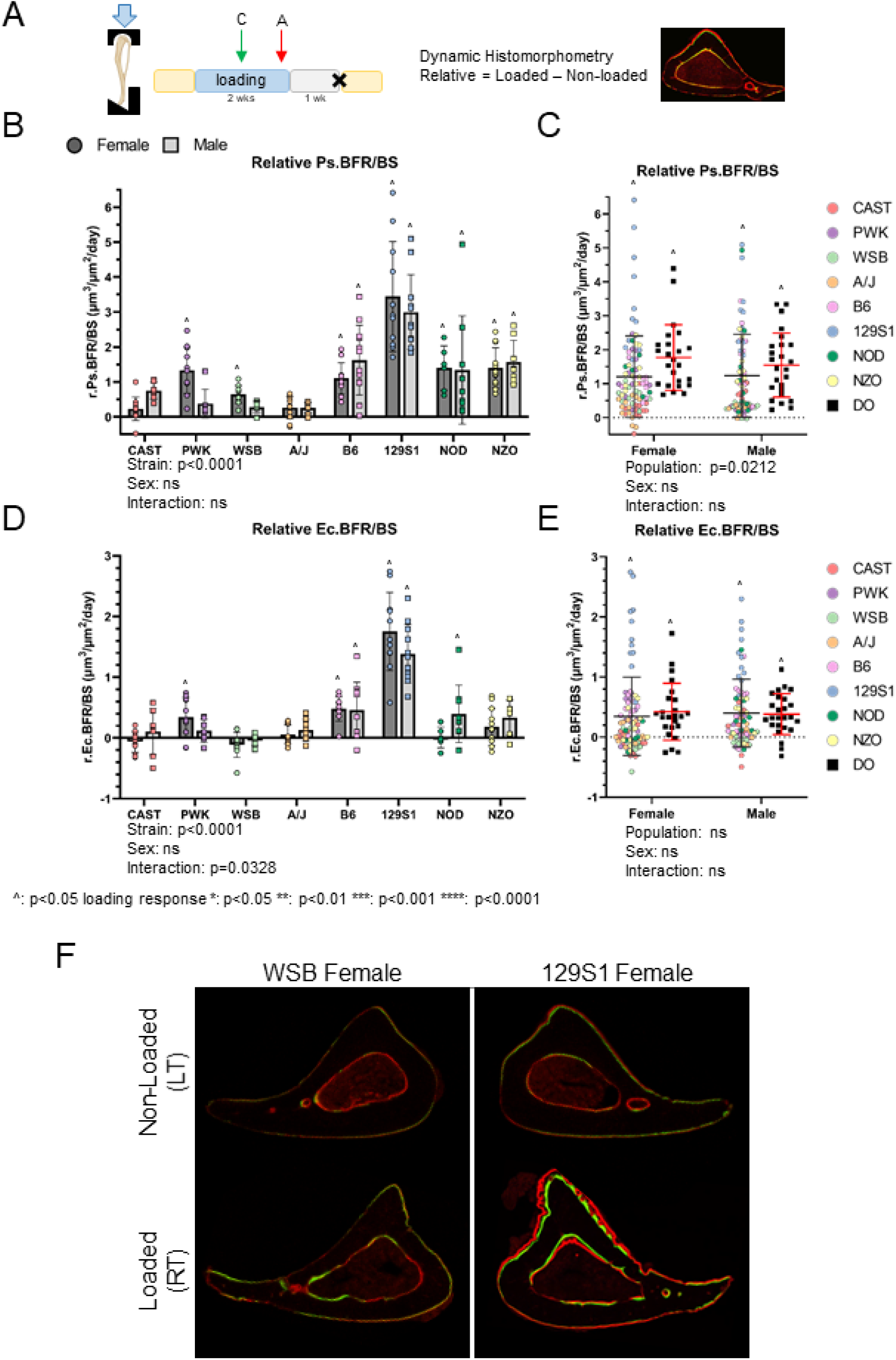
Loading-induced increases in bone formation rate vary with mouse strain. (A) Bone formation indices were measured on nonloaded and loaded tibia using classical dynamic histomorphometry. Relative changes were calculated with respect to nonloaded tibia. (B) Relative bone formation rate on the periosteal (Ps.) surface in eight inbred strains and (C) in inbred strains (pooled) and diversity outbred (DO) mice. (D) Relative bone formation rate on the endocortical (Ec.) surface of inbred strains and (E) in inbred strains (pooled) and DO mice. (F) Representative histomorphometry images of WSB and 129S1. Two-way ANOVA (sex and strain or population). Data shown as mean±SD.

On the endocortical surface, significant increases in bone formation indices were observed in fewer than half of the strain/sex groups, and the magnitudes of increase were less than on the periosteal surface. The effect of sex on endocortical indices was marginal, with the only pairwise difference between sexes in mineralizing surface in NOD mice. The response to loading did not differ between DO and Inbred Founder populations (Fig. 4E, Suppl. Fig. S6F,H).

In summary, we observed highly significant strain-dependent variation in 2D bone formation indices on periosteal and endocortical surfaces, with 129S1 mice again showing the greatest increases due to loading. The DO population had a greater periosteal loading response than the average of the Inbred Founders. Sex-dependent effects were marginal.

### Bone Response to Mechanical Loading is Highly Heritable

Broad-sense heritability of each loading response variable was calculated for Inbred Founder strains (Table 1). The five measures related to cortical bone size and the four related to periosteal bone formation were all highly heritable (H^2^ = 0.79-0.95), indicating that 79 to 95% of the variation in these traits is due to genetic factors. Several endocortical measures of bone formation also had high heritability, including Ec.BFR/BS, Ec.MAR and Ec.Form/BV (H^2^ > 0.92).

**Table 1:** Broad-sense heritability (H^2^) of all relative outcomes (relative = loaded – nonloaded) measured in the Inbred Founders.

| **Measurement criteria** | **Trait** | **Heritability (H<sup>2</sup>)** |
|---|---|---|
| µCT morphometry | Ct. Ar. | 0.95 |
|  | Ct.Th. | 0.95 |
|  | pMOI | 0.89 |
|  | Ct.Ar./Tt.Ar. | 0.86 |
|  | Tt.Ar. | 0.85 |
|  | Ma.Ar. | 0.55 |
|  | TMD | 0.46 |
| Dynamic histomorphometry | Ec.BFR/BS | 0.97 |
|  | Ps.MAR | 0.95 |
|  | Ps.BFR/BS | 0.94 |
|  | Ec.MAR | 0.94 |
|  | Ps.MS/BS | 0.87 |
|  | Ec.MS/BS | 0.32 |
| Timelapse µCT | Ec.Form/BV | 0.92 |
|  | Ps.Form/BV | 0.79 |
|  | Ec.Resorb/BV | 0.44 |
|  | Ps.Resorb/BV | 0.36 |

### Mouse Strains with More Elongated Osteocyte Lacunae Are More Responsive to Loading

Because homeostatic bone phenotypes (13) and responsiveness to loading are both heritable in Inbred Founder mice (Table 1), we examined correlations between loading response outcomes and baseline (pre-scan) measures of bone morphology. Loading response variables correlated only weakly (or not at all) with most measures of bone morphology in Inbred Founders (p < 0.05, R^2^ = 0.03-0.17,) and in DO mice (R^2^= 0.08-0.25, Suppl. Fig. S7A,B). The only exception was loading-induced change in polar moment of inertia (pMOI, a measure of resistance to bending and torsion), which was moderately correlated with baseline cortical area, total area, marrow area, and pMOI in Inbred mice (R^2^ = 0.27-0.44).

Because osteocytes are mechanosensitive and may regulate the bone response to loading (1, 23), we next examined the relationship of osteocyte lacunar morphology with loading response. We previously measured lacunar traits in tibial samples from the eight Inbred Founder strains using high-resolution x-ray microscopy (13). Correlation analysis between average values of these traits and average loading response outcomes from the 16 inbred strain/sex groups revealed many significant correlations (p < 0.05, R^2^ = 0.25 – 0.75; Fig. 5A). Notably, most of these correlations were substantially stronger than ones between loading responses and bone morphology traits (Suppl. Fig. S7). In particular, lacunar volume density (Lc.Vol/TV) positively correlated with many loading outcomes; indicating that inbred mice with greater volume fractions of lacunae were more responsive to loading (R^2^ = 0.48; Fig. 5B). Lacunar aspect ratio (Lc.AR) was strongly and negatively correlated with many loading outcomes, indicating that inbred mice with more elongated lacunae were more responsive to mechanical loading (R^2^ = 0.62; Fig. 5C). In summary, measures of osteocyte lacunar morphology are bone-intrinsic traits that strongly correlate with loading response across Inbred mice.

**Fig. 5:**
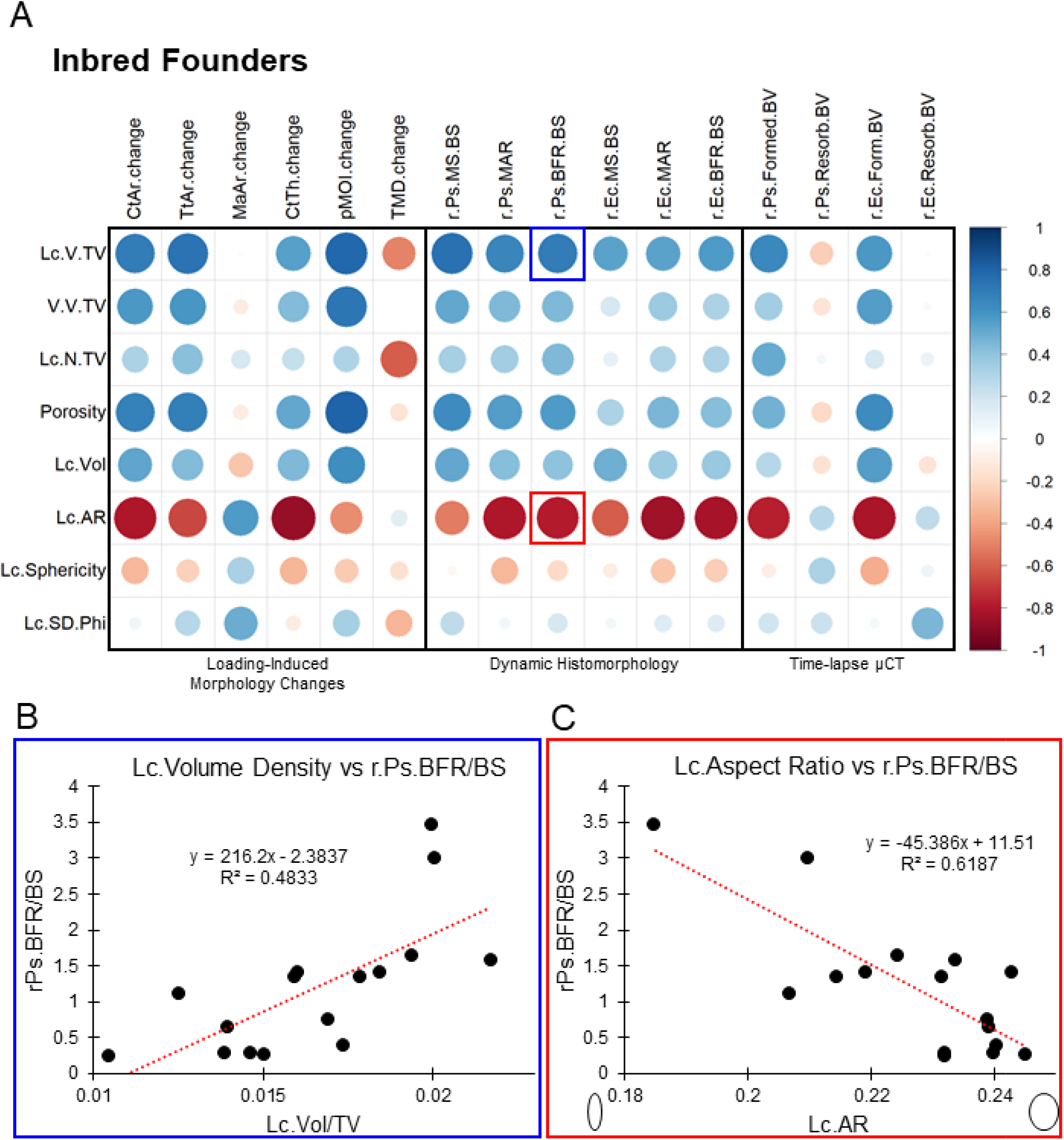
Osteocyte lacunar morphology correlates with loading response. Lacunar traits were measured in male and female mice from eight inbred strains using high-resolution X-ray microscopy (reported in Migotsky et al 2024). (A) Pearson’s correlation matrix (ρ values) based on average values for each strain/sex (n=16). Scalebar indicates correlation. (B) Average relative (loaded – nonloaded) periosteal bone formation rate (Ps.BFR/BS vs average lacunar (Lc.) aspect ratio. (C) Average relative Ps.BFR/BS vs average lacunar volume density.

### The Transcriptional Response to Loading is Largely Conserved Between High and Moderate Responder Strains

We next examined the bone transcriptional response to loading in high-responder 129S1 mice and compared it to our published dataset in age- and sex-matched B6 mice subjected to the same loading protocol (2); B6 mice are widely used in bone loading studies and were moderate responders here. Bulk RNAseq was performed on cortical bone tissue collected after one or five bouts of daily loading; differentially expressed genes (DEGs) were determined by comparison of paired loaded and non-loaded tibiae. After one bout of loading, 19 and 40 loading-related DEGs were identified in B6 and 129S1, respectively, with 13 shared DEGs (Fig. 6A). The genes showing the largest fold-changes with loading were largely the same in both strains, including *Wnt1* and *Ngf*, genes with known roles in loading-induced bone formation (28, 29) (Fig. 6B). Genes unique to 129S1 were *Ptgs2*, *Gpr3*, and *Egr2*. *Ptgs2* (*COX2*) is a well-known mechanoresponsive gene that can stimulate osteoblast differentiation (30), while *Egr2* is important to bone and nerve development (31).

**Fig. 6:**
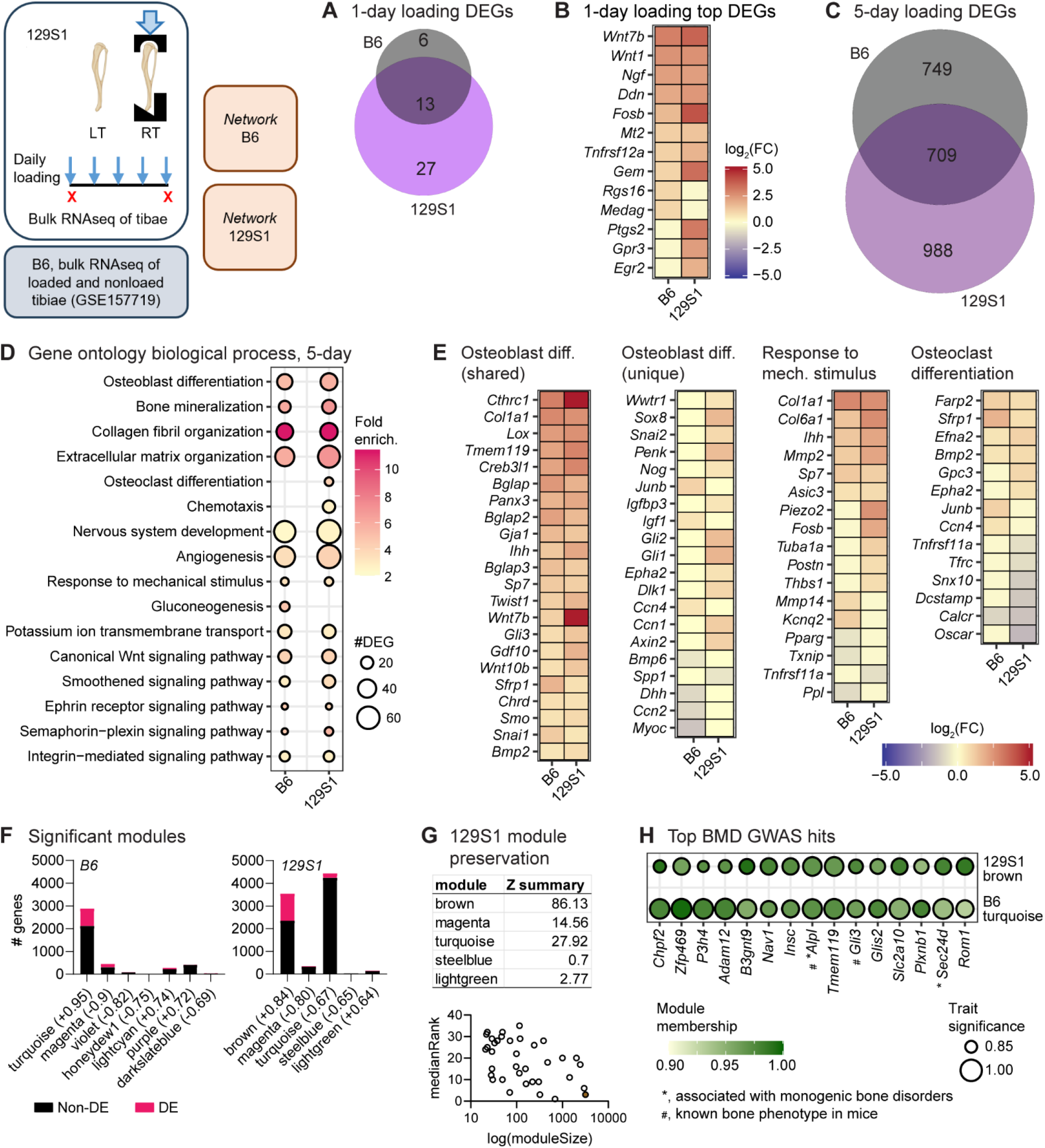
Loading response is largely conserved between B6 and 129S1 at transcriptome level. Female 129S1 mice (high responders) were subjected to five loading bouts and mRNA collected from cortical bone of bilateral tibias. RNAseq analysis was performed and compared with a published dataset from age- and sex-matched B6 mice (moderate responders, reported in Chermside-Scabbo et al 2020). (A) No. of shared and strain-specific DEGs between B6 and 129S1 following 1-day and (C) 5-day loading. (B) Expression of top DEGs following 1-day loading. (D) Gene Ontology biological processes enriched among DEGs from 5-day loading. (E) Expression of genes in selected loading-induced bone formation-related processes. (F) Significant (p<0.05) WGCNA modules and their correlation (in parenthesis) with loading. (G) Preservation score (Z_summary_) and median rank of 129S1 modules in B6. Filled circle denotes the highly preserved 129S1-brown module. (H) Top 15 shared hub genes associated with human BMD GWAS in loading-associated modules.

After 5 days of loading, we identified 1458 and 1697 DEGs in B6 and 129S1, respectively (Fig. 6C). GO analyses showed similar process enrichment in B6 and 129S1, including *osteoblast differentiation* and *bone mineralization*, as well as *canonical Wnt signaling*, a known regulator of bone’s anabolic response to loading (2, 28, 32) (Fig. 6D). Analysis of the 709 DEGs shared between 5-day B6 and 129S1 revealed enrichment of many of these same osteogenic processes – *bone mineralizatio*n, *osteoblast differentiation*, *BMP signaling*, and *canonical Wnt signaling pathway* (Suppl. Table S1). The unique DEGs in B6 (749) were enriched in *bone development*, *ECM organization*, and *cell population proliferation* (Suppl. Table S1). The 129S1-unique DEGs (988) demonstrated enrichment similar to those of the shared and B6-specific genes but also exhibited enrichment for *osteoclast differentiation* (Suppl. Table S1). Altogether, this analysis suggests that the transcriptional response to loading is largely conserved between B6 and 129S1 strains.

We next explored the expression levels of genes mapped to specific biological processes (Fig. 6E). In *osteoblast differentiation*, key genes such as *Tmem119*, *Bglap*, *Sp7*, and *Bmp2* were upregulated with loading in both strains, suggesting a shared activation of core pathways. Interestingly, in 129S1 loading upregulated *Snai2* and downregulated *Spp1* (O*steopontin*). *Snai2* maintains bone marrow stromal cells (BMSC) by suppressing *Spp1* expression, which inhibits self-renewal of BMSC (33, 34). This suggests that the loading response in 129S1 may involve additional progenitor cell regulatory mechanisms beyond those observed in B6. In addition, *Gli1* and *Gli2* were upregulated exclusively in 129S1, indicating a possible strain-specific enhancement of osteoblast differentiation related to hedgehog signaling (35). *Ccn1* was also upregulated only in 129S1; this gene is required for PTH-induced bone formation (36) and plays a key role in *Piezo1*-mediated osteocyte mechanotransduction (37). In the *Response to mechanical stimulus biological* process, *Piezo2* was upregulated with loading only in 129S1 (Fig. 6E). *Piezo2* has been shown to share some of the Piezo1 functions in osteoprogenitors, where combined deletion of *Piezo1* and *Piezo2* exacerbates bone deformities and/or spontaneous fractures compared to *Piezo1* loss alone (38). *Piezo2* transmits high strain-mediated Ca^2+^ signaling in chondrocytes and, and high strain also increases *Piezo2* expression (39, 40). Therefore, upregulation of *Piezo2* along with *Ccn1* may contribute to increased mechanosensitivity in 129S1 compared to B6.

Many genes related to osteoclast differentiation and bone resorption were uniquely differentially expressed in 129S1 (Fig. 6E and Fig. S8). Genes essential for osteoclast differentiation and function such as *Tfrc* (encodes *Tfr1*) and *Snx10* were downregulated with loading in 129S1 (41–43). Markers of mature multinucleated osteoclasts e.g., *Tnfrsf11a* (*Rank*), *Ctsk*, *Acp5*, *Calcr*, *Dcstamp*, *Ocstamp*, *Oscar*, and *Atp6v0d2*, were also downregulated in 129S1. While these positive regulators of osteoclast differentiation were downregulated, the negative regulators, *Cartpt* and *Tnfrsf11b* (*Opg*), were upregulated in 129S1. Overall, these differences in osteoclast genes suggest a stronger anti-resorptive transcriptional response to loading in 129S1.

We next identified coordinated gene expression modules associated with loading responses in B6 and 129S1. The modules with the strongest positive loading correlations were *turquoise* in B6 and *brown* in 129S1 (Fig. 6F). Strong preservation of 129S1 *brown* module was observed in B6 (Z_sum_ > 10, median rank = 3), which largely overlapped with B6 *turquoise*. This suggests that the gene co-expression networks associated with loading response are mostly conserved between the strains (Fig. 6G and S9A). B6 *turquoise* and 129S1 *brown* were strongly enriched for mouse homologs of human genes implicated by BMD GWAS (Suppl. Table S3 and Fig. S9B) (44). Among the top 15 shared GWAS genes (ranked by module kME), several have been implicated in monogenic bone disorders in humans and bone phenotypes in mice, including *Alpl*, *Sec24d*, and *Gli3* (Fig. 6H) (44). Additionally, we identified several genes with no previously described role in bone: *Zfp469*, *P3h4*, *B3gnt9*, *Nav1*, and *Insc*. In summary, transcriptional responses to loading appear to be largely conserved between B6 and 129S1, although upregulation of some mechanoresponsive genes and regulation of osteoclastic genes were unique to 129S1. Additionally, co-expression modules associated with loading are enriched for genes implicated by human BMD GWAS.

## Discussion

We employed two murine models of genetic diversity to assess the influence of genetic background on the skeletal response to mechanical loading. Our results revealed that bone’s response to mechanical loading is highly heritable. Measures of bone formation due to loading varied significantly among inbred mouse strains, and we also observed strain-sex interactions across several measurements, with certain strains showing pronounced sexual dimorphism. A cohort of DO mice, a relatively new model of genetic diversity, showed a response to loading that was higher than the average of the inbred strains. Furthermore, we demonstrated that some osteocyte lacunar morphology traits correlated with the magnitude of the loading response across inbred strains. Remarkably, 129S1, the strain with the most elongated lacunae exhibited the highest magnitude of response on both periosteal and endocortical surfaces. 129S1 also demonstrated a robust transcriptional response to loading. Overall, genetic background significantly contributes to cortical bone response to loading, indicating genetic control of mechanoresponsiveness.

Many skeletal traits are highly heritable. In humans, heritability of BMD is estimated to be between 0.5 and 0.85 (45), and susceptibility to osteoporotic fracture is also reported to be heritable (46, 47). In addition, bone morphological traits in humans and mice are highly heritable (48–51). We recently reported that bone mass and many morphology and mechanical traits are heritable across eight inbred mouse strains that account for ∼90% of genomic variation in *M. musculus* (13). Responses to exercise in both humans and mice also exhibit heritable variation (52, 53). Kesavan et al. reported that bone loading responses were heritable in B6×C3H F2 mice (10, 54). Our current study provides new evidence, incorporating greater genetic variation and more allele combinations, that loading-induced changes in bone morphology and bone formation measures have heritability estimates ranging from 45% to 97%, supporting the conclusion that response to loading is strongly influenced by genetic background.

Inbred 129S1 mice demonstrated the highest responsiveness to loading across the eight inbred strains. Conversely, A/J and WSB exhibited the lowest response, while B6 displayed a moderate response. WSB and A/J have been shown to lose trabecular bone mass due to unloading (14, 15), suggesting these strains are sensitive to unloading despite being relatively unresponsive to tibial (over)loading. By contrast, 129S1 and B6 respond bidirectionally; while they lose trabecular bone mass with unloading (14, 15), in the current study they gained cortical bone with loading. Notably, 129S1 females showed an almost 6-fold increase in periosteal mineral apposition rate with loading in the present study, inclusive of both lamellar and woven bone formation. We observed significant bone formation on the endocortical surface as well. This heightened response in 129S1 may be due to mechanical factors, intrinsic biological factors, or both. Mechanically, we applied equivalent tensile strain at the anteromedial surface, although the compressive strain at the posterolateral surface might differ between mouse strains due to differences in tibial geometry. Biologically, we observed greater fold changes in mechanoresponsive genes such as *Wnt1* and *Wnt7b*, consistent with a heightened mechanosensitive transcriptional response in 129S1 compared to B6. However, because B6 and 129S1 DEGs were derived from independent studies, direct comparisons should be interpreted with caution. Nonetheless, some mechanoresponsive genes (*Piezo2*, *Ptgs2*, and *Thbs1*) were upregulated only in 129S1, further supporting heightened mechanosensitivity in this strain. In addition, 129S1 exhibited a larger number of differentially expressed osteoanabolic genes than B6, which may explain the increased osteoblast activity marked by the woven bone. While network module preservation statistics showed that core transcriptional responses were conserved between strains, 129S1-unique genes alone were sufficient to robustly enrich several processes, including *osteoblast differentiation*, *BMP signaling* and *smoothened signaling*. The lower expression of osteoclast-associated genes in 129S1 was also intriguing, as it suggests a depletion of osteoclast numbers and/or activity on the endocortical surface. Future studies might use 129S1 to further explore this phenomenon.

Baseline differences in morphology and transcriptomes among laboratory mouse strains can influence loading response. For example, A/J mouse bones have greater baseline expression of Wnt pathway inhibitory genes (55); this may explain why they did not respond to loading in our study. Interestingly, in the present study the low-responder WSB females exhibited a relatively high (>60%) Ps.MS/BS at baseline, which increased to >80% following loading. Thus, the relative change in Ps.BFR/BS was modest because baseline activity was already high. Future studies might explore the basis for baseline differences in osteoblast activity across inbred strains.

DO mice exhibited variance in their loading responses, with a range similar to that of the eight inbred strains; this is consistent with each DO mouse having a unique mixture of alleles of the inbred founder strains. However, on average the DO mice had a greater response to loading than the founders. We observed significantly greater increases in total area and pMOI, and most loading outcomes were higher in the DO population than the pooled Inbred Founder population. This may be attributed to epistatic regulation, in which a gene’s phenotypic effect depends on its interaction with other genes rather than acting independently (56). As a result, a particular combination of gene variants can produce a phenotype that is muted once that exact combination is disrupted, even if each variant is still present individually. We observed similar findings previously while phenotyping these mice, where outbreeding eliminated extreme phenotypes seen in the founders (13). The genome-wide heterogeneity of DO mice, which approaches that of humans, combined with the ability to perform multi-scale phenotyping, has made them a valuable complement to human genetic studies. For example, sequencing of DO mouse bones and cells were used to identify novel driver genes underlying human BMD genome-wide association study (GWAS) associations (17, 18). Al-Barghouthi et al. used gene expression data from bone tissue in DO mice to build co-expression networks and identify novel candidate genes underlying human BMD associations, and performed GWAS in DO mice for multiple bone traits, including bone strength(17). Dillard et al. applied single-cell sequencing to determine not only which genes are likely responsible for BMD associations, but also in which bone cell types and stages of differentiation they act (18). Together, these studies reinforce the utility of DO mice for investigating therapeutic interventions and systems genetics through their capture of natural variation.

Over the past three decades, multiple groups have published studies examining the role of mechanical loading in rodents (2, 3, 57–59). However, only a limited number of these studies have investigated sexual dimorphism by including both female and male animals (7, 21, 22, 60, 61). In the present study, we examined both sexes in all inbred strains and DO mice. Our data reveal strong strain-sex interactions across several measurements, including periosteal and endocortical bone formation rates, suggesting that sex may modulate mouse strain-specific responses to mechanical loading. This pattern is consistent with the baseline bone traits we previously reported for these inbred strains (13). The sex effect, however, was more pronounced in some strains than others, with the largest differences observed in the wild-derived CAST and PWK strains, as well as the 129S1 strain. Specifically, PWK males showed almost no response to loading, while the females demonstrated a modest response. This discrepancy further highlights the importance of including sex as a biological variable in such studies. Nonetheless, when all inbred founder strains were pooled to analyze as a population, no overall sex-based difference in response to mechanical loading was detected. Moreover, no significant differences were observed between DO males and females. Collectively, these findings indicate that the influence of sex is strain-dependent and secondary to the influence of genetic background. We note that these results are based on WT mice from the populations we studied and may not translate to effects in mutant mice. Inclusion of sex as a biological variable remains an important consideration in the design of loading studies.

We observed some significant correlations between osteocyte lacunar morphology and loading response in inbred mice. Strains with higher lacunar volume density and more elongated lacunae responded more to mechanical loading. Since osteocytes are the primary mechanosensitive cells in the bone tissue, it is perhaps unsurprising that lacunar volume density may influence bone formation. By contrast, we did not observe a strong correlation between lacunar number density (Lc.N/TV) and bone formation (R^2^=0.26) (Fig. 5A). This is consistent with findings from Robling and Turner who examined lacunar number density in B6, C3H, and DBA inbred mice in relation to their response to axial loading and found no correlation between lacunar density and mechanosensitivity (9). Notably, we identified lacunar shape as the strongest correlate with loading response. The highest responder 129S1 had the most elongated lacunae along the loading axis, and the correlation across eight inbred strains indicates that more elongated lacunae are associated with a greater response to loading. This finding is supported by a recent study which used fluid-structure interaction modeling to show that elongated lacunae amplify interstitial fluid velocity and osteocyte strain (27). Together, our findings and those of others suggest that lacunar shape is a strong modulator of osteocyte mechanosensation (62, 63).

Our network analysis in B6 and 129S1 shows that loading-correlated gene sets are enriched for mouse homologs of human genes implicated by BMD GWAS. Several of these genes were highly connected module members (high kME), meaning that their expression closely tracked the module eigengene following loading. Some GWAS genes were unique to either 129S1 or B6. Functional studies of the genes unique to 129S1 may help reveal the biological basis for 129S1’s heightened mechanoresponsiveness (Suppl. Fig. S9C). Together, these findings suggest: 1) many mechanosensitive genes are related to bone disorders, highlighting the importance of mechanoresponsiveness in bone health; and 2) while some highly connected module members have established roles in bone biology in mice or humans, many others have not yet been functionally linked to bone, such as *Zfp469*, *P3h4*, *B3gnt9*, *Nav1*, and *Insc*. These module members represent promising candidates for further investigation in bone pathogenesis and therapeutic development.

There are limitations to our study. First, differences in baseline activity among mouse strains could alter the response to our specific loading intervention. To control for this, all loading parameters were reported as relative values in the loaded limb compared to the non-loaded limb. Second, the estimation of loading force for DO mice has a large range of errors. While the mean error of the model was 32 µε, the range was approx. ± 500 µε. However, this error was distributed around zero across the range of loading forces (Suppl. Fig S2A) and there was no correlation between the estimated strain produced by the loading force and the loading response (Suppl. Fig S2B). Thus, while the error of the force estimation adds variability to the population it did not bias our results. Third, the values for the lacunar traits used in our correlation were not measured from the same animals that were loaded, but were average values for each strain/sex group reported in Migotsky et al (13). Although the inbred mice within a strain/sex group are genetically identical between the two studies, there could be differences due to batch effects. Fourth, because the B6 and 129S1 RNAseq samples were processed in different batches, fold changes for individual genes are not directly comparable. Despite these limitations, this study also has several strengths. Our large sample size increased the statistical power, reduced the influence of individual variability, and improved confidence in the observed outcomes. Additionally, we incorporated broad genetic diversity spanning eight inbred strains and the DO mice. We have also developed a method to load genetically unique animals without the need for strain gauging, illustrating an approach for future use of DO mice to study bone mechanobiology (Fig. S1). Finally, we used multiple measures (bone morphometry, time-lapse analysis, dynamic histomorphometry) that support the same overall conclusion that loading responses are highly heritable.

Taken together, our findings demonstrate the substantial impact of genetic background on the cortical bone response to mechanical loading, establishing the role of genetics in determining mechanoresponsiveness across a diverse population. The response to mechanical loading, measured by multiple methods, significantly varied among inbred mouse strains and correlated with lacunar morphology. We showed an increased response to loading in the outbred population compared to the Inbred Founders. Furthermore, we identified the inbred 129S1 strain as a model of heightened mechanosensitivity that shares transcriptional similarity with the lab-standard B6 strain, making it useful for future studies of bone mechanobiology. Additionally, this work provides groundwork and rationale to perform GWAS and quantitative trait locus (QTL) mapping to identify candidate genes regulating the response to loading in DO mice. Uncovering novel genes can provide new targets for therapeutics to increase bone mass, especially in individuals incapable of mechanical loading, exercise, or with reduced mechanosensation.

## Methods

This study was approved by the Washington University Institutional Care and Use Committee (IACUC; protocol #22-0384) and performed at Washington University in St. Louis, St. Louis, MO). Experimental plan is shown in Fig. 1.

### Mice

To model genetic diversity, two mouse populations were utilized: (1) eight Inbred Founder strains: CAST/EiJ (JAX Stock #000928), PWK/PhJ (#003715), WSB/EiJ (#001145), A/J (#000646), C57BL/6J (#000664), 129S1/SvImJ (#002448), NOD/ShiLtJ (#001976), and NZO/HlLtJ (#002105) and (2) Diversity Outbred (DO) mice (#009376) (Fig. 1A). Females and males were included in equal numbers. Inbred mice (n = 12/strain/sex) were received at 8 weeks age from JAX and aged to 22 weeks in our animal facilities following standard housing protocol. Due to attrition (see Suppl. Methods), a total of N = 169 Inbred Founder mice were analyzed at the end of the study. The DO mice (n = 25/sex) were obtained between 3 and 5 weeks age and similarly aged to 22 weeks. One female DO mouse died during µCT; a total of 49 DO mice were analyzed at the end of the study.

### Axial tibial loading

Mice were loaded in axial tibial compression for 2 weeks to stimulate cortical bone formation (Fig. 1B). While under anesthesia (Isoflurane 2-4% v/v) the right tibia was placed in a custom-built holder and loaded to a peak axial force to engender an approximate tensile strain of 1500 με (see *Calculations of Necessary Force* below) using a dynamic testing machine (ElectroPuls 1000, Instron) as previously described (57, 64). The left tibia remained non-loaded as an internal, contralateral control. Mice were loaded using a haversine waveform for 60 cycles per day at 4 Hz, 5 days per week for 2 weeks. One week after the last bout of loading, mice were euthanized by CO_2_ asphyxiation.

### Calculations of necessary force for strain-matching

All mice were loaded to an estimated equivalent peak tensile strain of 1500 με on the antero-medial surface (i.e., strain-matched). To calculate the force necessary to engender 1500 με, a calibration set of non-loaded Inbred Founder strains were strain gauged (mice used in Migotsky et al. (13)). Briefly, the antero-medial surface of the right tibia was exposed *in situ* and a uniaxial strain gauge was glued 5 mm proximal to the TFJ (approx. 37% of tibial length (64); Suppl. Fig. S1A). The tibia was loaded from 2–12 N while force and mechanical strain were recorded. From the force-strain linear regression, the force corresponding to 1500 με was calculated per animal (Suppl. Fig. S1B). An average value per strain/sex group was calculated from n = 4-9 mice (Suppl. Fig. S1C).

Because each DO mouse is genetically unique, strain gauging is not a feasible method to calculate the necessary loading force per animal. Therefore, we used the tibial strain gauge and morphology data from the above Inbred Founder mice (N = 130 animals) to build a multivariable linear regression to predict necessary loading force using non-invasively measured traits from μCT. Starting with 11 traits (Ct.th, TMD, Tt.Ar, Ct.Ar, pMOI, weight, AreaRatio, Circularity, Eccentricity, Extent Inside, Perm Ratio Ellipse), backward elimination was used to remove non-significantly contributing variables, leaving five variables in the linear regression: Tt.Ar, Ct.Ar, weight, AreaRatio, and Eccentricity. The regression explains 57% of the variation in the population (Suppl. Fig. S1D,E). To validate the accuracy of the model, strain gauging was performed on the left, non-loaded tibia of DO mice (n = 16-17 per sex) postmortem. The error between loading force based on the model and measured force was randomly distributed around zero (Suppl. Fig. S2A). The mean error in measured strain was 32 με (range -437 to +499 με) and did not correlate with bone formation response of the contralateral limb, indicating no bias (Suppl. Fig. S2B).

### MicroCT (µCT)

Serial microCT scanning was used to evaluate changes in the morphology of the tibia before and after loading. One to two days prior to loading, the left and right tibiae were scanned *in vivo* (referred to as pre-scan) at 10.5 μm/pixel resolution 5 mm proximal to the TFJ (vivaCT 40, Scanco Medical, -70 kVP, 8 W, 300 ms integration time, 213 slices). The same region was scanned *in situ* 1 week after the last day of loading (post-euthanasia, referred to as post-scan) using the same settings (Fig. 1B). Loading-induced changes in morphology per animal were calculated as the change in the loaded bone over time minus the change in the non-loaded bone over time (ΔΔ_µCT_ = (Post_loaded_-Pre_loaded_) -(Post_non-loaded_-Pre_non-loaded_)). To calculate loading-induced formed and resorbed bone volumes we performed time-lapse µCT analysis following Birkhold et al (65), and describe in detail elsewhere (66). In brief, pre- and post-scans of the same limb were registered in 3D and resampled to the same coordinate system. Formed volumes were defined as voxels present in the post-scan but not the pre-scan. Resorbed volumes were defined as voxels present in the pre-scan but not the post-scan (Suppl. Fig. S3A). The periosteal and endocortical surfaces were analyzed separately. Formed and resorbed volumes were normalized to the original cortical bone area. Relative values for all outcomes were calculated as loaded minus non-loaded.

### Dynamic histomorphometry

Calcein green (10 mg/kg)and alizarin red (30 mg/kg) were injected at the end of the first and second weeks of loading, respectively, to allow for visualization of newly mineralized bone. After the µCT post-scan, the left and right tibiae were collected and fluorochrome-labeled surfaces were analyzed (Osteo II, Bioquant) separately on the periosteal and endocortical surfaces (67). Relative values for bone formation indices were calculated as loaded minus non-loaded. Further details are provided in the Supplementary Methods.

### Heritability calculations

The broad-sense heritability (H^2^) of all relative loading responses in the Inbred Founder mice was determined to assess the proportion of variance attributed to genetic differences, following established methodology (68). Heritability was calculated using the formula H^2^ = σ^2^_strain_/(σ^2^_strain_+ σ^2^_sex_+ σ^2^_res_), where σ^2^_strain_ denotes the variance between strains, σ^2^_sex_ denotes the variance between sexes, and σ^2^_res_ denotes the residual variance. The variances were computed based on the sum of squares (SS) derived from a two-factor analysis of variance (ANOVA) defining strain and sex as the factors. Specifically, σ^2^_strain_ was calculated as SS_strain_/n_avg_ (where n_avg_ corresponds to the average group size), σ^2^_sex_ was determined as SS_sex_/df_sex_, (where σ^2^_res_ corresponds to SS_res_/df_res_).

### Correlation determination

To examine the relationship between bone morphology and bone response to loading, we performed several analyses. First, a correlation matrix of Pearson’s r coefficients was computed to assess the association between all initial μCT outcomes of the right tibia and the loading response outcomes per animal in both the Inbred Founder and DO populations. Second, a correlation matrix of Pearson’s r coefficients was calculated to examine the correlation between average values of osteocyte lacunar traits and the average loading responses per strain/sex group within the Inbred Founder population. The average lacunar trait values were previously published in Migotsky et al. (13). Bivariate plots were also generated to visualize the relationships between lacunar traits and loading responses.

### Bulk RNA sequencing of bone tissue

Based on preliminary results of bone formation outcomes in Inbred Founders, we identified female 129S1 as the highest responding strain and performed RNAseq on a separate cohort of these mice. 22-week-old female 129S1 mice were subjected to unilateral tibial loading (as described above) for 1 or 5 consecutive days (n = 3 and 5 mice/timepoint, respectively). Mice were euthanized 4 h after the last loading session, and total RNA was extracted from cortical bone tissue as described (2). Sequencing was performed on an Illumina NovaSeq 6000. DESeq2 in R/Bioconductor with Wald test was used to determine the differentially expressed genes (DEGs) between loaded and non-loaded conditions, with nonloaded as baseline. To compare these results with B6 loading response, we used a dataset published by our group (GSE157719, utilized C57Bl/6N mice) which has a similar experimental design as the current study (2). Gene Ontology (GO) enrichment analysis was performed using the database for annotation, visualization, and integrated discovery (DAVID) (69, 70). Further details are included in the Supplementary Methods.

### Co-expression network analysis

Weighted gene co-expression network analysis (WGCNA) was performed on 5-day 129S1 samples from the current study and the historical 5-day B6 samples separately to identify modules of co-expressed genes (n=5/group) (71). The expression profile was then correlated with loading condition (loaded or nonloaded) within each network. Module preservation was tested to assess how well network structure of 129S1 modules were preserved in the B6 dataset (72). To identify modules of genes enriched for BMD GWAS genes, we obtained the mouse homolog list curated by Sabik et al (44) and used Fisher’s exact test to measure the overrepresentation of GWAS genes in each module of 129S1 and B6 separately. Benjamini-Hochberg correction was then applied to correct for multiple testing. More details are provided in Supplementary Methods.

### Statistical analysis

Statistical analysis was conducted using GraphPad Prism (v9). Relative loading response outcomes from the eight Inbred Founders were analyzed using a two-factor analysis of variance (ANOVA) with mouse strain and sex as factors. Similarly, outcomes from the Inbred Founder population and the DO population were compared using a two-factor ANOVA with population and sex as factors. Post-hoc tests were conducted with Sidak correction. To evaluate the significance of the loading response within a given strain/sex group, the value of the right tibia (loaded) was compared to the value of the left tibia (non-loaded) during post-hoc testing. Significance was set at p < 0.05. Pearson’s correlation coefficients were calculated using the *cor* function in R (v4.0.2), and the resulting correlations were visualized using the *corrplot* function. All bioinformatics were performed using R (v4.4.3) and Bioconductor (v3.20).

## Acknowledgments

We thank the following for their support in this project: WashU Musculoskeletal Research Center (MRC -NIH P30 AR074992) for support of microCT and mechanical loading resources; WashU Genome Technology Access Center for sequencing services; NIH/ NIAMS R01 AR047867, NIH T32 DK108742, NIH T32 EB018266, NIH R21 079052, NIH S10 RR023660, and Jax DO Grants.

## References

1. A. G. Robling, A. B. Castillo, C. H. Turner, Biomechanical and molecular regulation of bone remodeling. Annu Rev Biomed Eng 8, 455–498 (2006).

2. C. J. Chermside-Scabbo, et al., Old Mice Have Less Transcriptional Activation But Similar Periosteal Cell Proliferation Compared to Young-Adult Mice in Response to in vivo Mechanical Loading. J Bone Miner Res 35, 1751–1764 (2020).

3. C. Chlebek, J. A. Moore, F. P. Ross, M. C. H. van der Meulen, Molecular Identification of Spatially Distinct Anabolic Responses to Mechanical Loading in Murine Cortical Bone. J Bone Miner Res 37, 2277–2287 (2022).

4. C. H. Turner, Functional determinants of bone structure: beyond Wolff’s law of bone transformation. Bone 13, 403–409 (1992).

5. J. Fritton, E. Myers, T. Wright, M. Vandermeulen, Loading induces site-specific increases in mineral content assessed by microcomputed tomography of the mouse tibia. Bone 36, 1030–1038 (2005).

6. S. J. Warden, I. S. Davis, M. Fredericson, Management and prevention of bone stress injuries in long-distance runners. J Orthop Sports Phys Ther 44, 749–765 (2014).

7. A. K. Nepal, et al., Methodological aspects of in vivo axial loading in rodents: a systematic review. J Musculoskelet Neuronal Interact 23, 236–262 (2023).

8. M. P. Akhter, D. M. Cullen, E. A. Pedersen, D. B. Kimmel, R. R. Recker, Bone response to in vivo mechanical loading in two breeds of mice. Calcif Tissue Int 63, 442–449 (1998).

9. A. G. Robling, C. H. Turner, Mechanotransduction in bone: genetic effects on mechanosensitivity in mice. Bone 31, 562–569 (2002).

10. C. Kesavan, et al., Identification of genetic loci that regulate bone adaptive response to mechanical loading in C57BL/6J and C3H/HeJ mice intercross. Bone 39, 634–643 (2006).

11. H. Yang, et al., Subspecific origin and haplotype diversity in the laboratory mouse. Nat Genet 43, 648–655 (2011).

12. A. Roberts, F. Pardo-Manuel de Villena, W. Wang, L. McMillan, D. W. Threadgill, The polymorphism architecture of mouse genetic resources elucidated using genome-wide resequencing data: implications for QTL discovery and systems genetics. Mamm Genome 18, 473–481 (2007).

13. N. Migotsky, et al., Multi-scale cortical bone traits vary in females and males from two mouse models of genetic diversity. JBMR Plus 8, ziae019 (2024).

14. M. A. Friedman, et al., Genetic variability affects the skeletal response to immobilization in founder strains of the diversity outbred mouse population. Bone Rep 15, 101140 (2021).

15. M. A. Friedman, et al., Genetic variation influences the skeletal response to hindlimb unloading in the eight founder strains of the diversity outbred mouse population. Journal Orthopaedic Research 42, 134–140 (2024).

16. K. L. Svenson, et al., High-resolution genetic mapping using the Mouse Diversity outbred population. Genetics 190, 437–447 (2012).

17. B. M. Al-Barghouthi, et al., Systems genetics in diversity outbred mice inform BMD GWAS and identify determinants of bone strength. Nat Commun 12, 3408 (2021).

18. L. J. Dillard, G. Calabrese, L. Mesner, C. Farber, Cell type-specific network analysis in Diversity Outbred mice identifies genes potentially responsible for human bone mineral density GWAS associations. Elife 13, RP100832 (2026).

19. L. I. Plotkin, A. Bruzzaniti, R. Pianeta, Sexual Dimorphism in the Musculoskeletal System: Sex Hormones and Beyond. J Endocr Soc 8, bvae153 (2024).

20. Y.-Y. Zhang, et al., Insights and implications of sexual dimorphism in osteoporosis. Bone Res 12, 8 (2024).

21. L. B. Meakin, G. L. Galea, T. Sugiyama, L. E. Lanyon, J. S. Price, Age-related impairment of bones’ adaptive response to loading in mice is associated with sex-related deficiencies in osteoblasts but no change in osteocytes. J Bone Miner Res 29, 1859–1871 (2014).

22. A. M. Rooney, et al., Bone mass and adaptation to mechanical loading are sexually dimorphic in adult osteoblast-specific ERα knockout mice. Bone 158, 116349 (2022).

23. L. F. Bonewald, M. L. Johnson, Osteocytes, mechanosensing and Wnt signaling. Bone 42, 606–615 (2008).

24. Y. Han, S. C. Cowin, M. B. Schaffler, S. Weinbaum, Mechanotransduction and strain amplification in osteocyte cell processes. Proc Natl Acad Sci U S A 101, 16689–16694 (2004).

25. C. Price, X. Zhou, W. Li, L. Wang, Real-time measurement of solute transport within the lacunar-canalicular system of mechanically loaded bone: direct evidence for load-induced fluid flow. J Bone Miner Res 26, 277–285 (2011).

26. A. Gupta, S. Saha, A. Das, A. Roy Chowdhury, Evaluating the influence on osteocyte mechanobiology within the lacunar-canalicular system for varying lacunar equancy and perilacunar elasticity: A multiscale fluid-structure interaction analysis. J Mech Behav Biomed Mater 160, 106767 (2024).

27. A. Muñoz, A. De Paolis, L. Cardoso, A. Carriero, Osteocyte-lacuna shape and canaliculi architecture dictate fluid flow around osteocyte, and strain of cell and bone matrix: implications for cell mechanobiology and bone fragility. Bone 200, 117613 (2025).

28. L. Y. Lawson, et al., LOADING-INDUCED bone formation is mediated by *Wnt1* induction in OSTEOBLAST-LINEAGE cells. The FASEB Journal 36, e22502 (2022).

29. I. Rajpar, et al., Osteoblast-derived Nerve Growth Factor is Required for Skeletal Adaptation to Mechanical Load and the Osteoanabolic Effect of Gambogic Amide in Mice. [Preprint] (2025).

30. X. Zhang, et al., Cyclooxygenase-2 regulates mesenchymal cell differentiation into the osteoblast lineage and is critically involved in bone repair. J. Clin. Invest. 109, 1405–1415 (2002).

31. A. Chandra, S. Lan, J. Zhu, V. A. Siclari, L. Qin, Epidermal growth factor receptor (EGFR) signaling promotes proliferation and survival in osteoprogenitors by increasing early growth response 2 (EGR2) expression. J Biol Chem 288, 20488–20498 (2013).

32. K. Sawakami, et al., The Wnt co-receptor LRP5 is essential for skeletal mechanotransduction but not for the anabolic bone response to parathyroid hormone treatment. J Biol Chem 281, 23698–23711 (2006).

33. Y. Tang, S. J. Weiss, Snail/Slug-YAP/TAZ complexes cooperatively regulate mesenchymal stem cell function and bone formation. Cell Cycle 16, 399–405 (2017).

34. Q. Wei, et al., Snai2 Maintains Bone Marrow Niche Cells by Repressing Osteopontin Expression. Developmental Cell 53, 503–513.e5 (2020).

35. H. Hojo, et al., Gli1 Protein Participates in Hedgehog-mediated Specification of Osteoblast Lineage during Endochondral Ossification. Journal of Biological Chemistry 287, 17860– 17869 (2012).

36. G. Zhao, et al., CCN1/Cyr61 Is Required in Osteoblasts for Responsiveness to the Anabolic Activity of PTH. Journal of Bone and Mineral Research 35, 2289–2300 (2020).

37. Y. J. Hu, et al., Piezo1-mediated mechanotransduction controls osteocyte maturation and dendrite development via a YAP-CCN-Src signaling axis. Nat Commun 16, 10859 (2025).

38. L. Qin, et al., Roles of mechanosensitive channel Piezo1/2 proteins in skeleton and other tissues. Bone Res 9, 44 (2021).

39. G. Du, et al., Roles of TRPV4 and piezo channels in stretch-evoked Ca^2+^ response in chondrocytes. Exp Biol Med (Maywood) 245, 180–189 (2020).

40. W. Lee, et al., Synergy between Piezo1 and Piezo2 channels confers high-strain mechanosensitivity to articular cartilage. Proc. Natl. Acad. Sci. U.S.A. 111 (2014).

41. B. K. Das, et al., Transferrin receptor 1-mediated iron uptake regulates bone mass in mice via osteoclast mitochondria and cytoskeleton. eLife 11, e73539 (2022).

42. E.-L. Stattin, et al., SNX10 gene mutation leading to osteopetrosis with dysfunctional osteoclasts. Sci Rep 7, 3012 (2017).

43. C. H. Zhu, L. R. Morse, R. A. Battaglino, SNX10 is required for osteoclast formation and resorption activity. J of Cellular Biochemistry 113, 1608–1615 (2012).

44. O. L. Sabik, G. M. Calabrese, E. Taleghani, C. L. Ackert-Bicknell, C. R. Farber, Identification of a Core Module for Bone Mineral Density through the Integration of a Co-expression Network and GWAS Data. Cell Rep 32, 108145 (2020).

45. J. B. Richards, H.-F. Zheng, T. D. Spector, Genetics of osteoporosis from genome-wide association studies: advances and challenges. Nat Rev Genet 13, 576–588 (2012).

46. J. A. Kanis, et al., A family history of fracture and fracture risk: a meta-analysis. Bone 35, 1029–1037 (2004).

47. Q. Wu, J. Jung, Genome-wide polygenic risk score for major osteoporotic fractures in postmenopausal women using associated single nucleotide polymorphisms. J Transl Med 21, 127 (2023).

48. D. Karasik, et al., Heritability and Genetic Correlations for Bone Microarchitecture: The Framingham Study Families. J Bone Miner Res 32, 106–114 (2017).

49. S. Demissie, et al., Proximal hip geometry is linked to several chromosomal regions: genome-wide linkage results from the Framingham Osteoporosis Study. Bone 40, 743–750 (2007).

50. W. G. Beamer, L. R. Donahue, C. J. Rosen, D. J. Baylink, Genetic variability in adult bone density among inbred strains of mice. Bone 18, 397–403 (1996).

51. C. H. Turner, et al., Genetic regulation of cortical and trabecular bone strength and microstructure in inbred strains of mice. J Bone Miner Res 15, 1126–1131 (2000).

52. R. Simonen, E. Levälahti, J. Kaprio, T. Videman, M. C. Battié, Multivariate genetic analysis of lifetime exercise and environmental factors. Med Sci Sports Exerc 36, 1559–1566 (2004).

53. M. P. Massett, B. C. Berk, Strain-dependent differences in responses to exercise training in inbred and hybrid mice. Am J Physiol Regul Integr Comp Physiol 288, R1006–1013 (2005).

54. C. Kesavan, D. J. Baylink, S. Kapoor, S. Mohan, Novel loci regulating bone anabolic response to loading: expression QTL analysis in C57BL/6JXC3H/HeJ mice cross. Bone 41, 223–230 (2007).

55. S. H. Schlecht, et al., Canalization Leads to Similar Whole Bone Mechanical Function at Maturity in Two Inbred Strains of Mice. J Bone Miner Res 32, 1002–1013 (2017).

56. J. D. Riordan, J. H. Nadeau, From Peas to Disease: Modifier Genes, Network Resilience, and the Genetics of Health. Am J Hum Genet 101, 177–191 (2017).

57. D. Sun, M. D. Brodt, H. M. Zannit, N. Holguin, M. J. Silva, Evaluation of loading parameters for murine axial tibial loading: Stimulating cortical bone formation while reducing loading duration. J Orthop Res 36, 682–691 (2018).

58. F. M. Lambers, F. A. Schulte, G. Kuhn, D. J. Webster, R. Müller, Mouse tail vertebrae adapt to cyclic mechanical loading by increasing bone formation rate and decreasing bone resorption rate as shown by time-lapsed in vivo imaging of dynamic bone morphometry. Bone 49, 1340–1350 (2011).

59. Y.-F. Hsieh, C. H. Turner, Effects of Loading Frequency on Mechanically Induced Bone Formation. Journal of Bone and Mineral Research 16, 918–924 (2001).

60. L. Y. Lawson, et al., Loading-Induced Bone Formation is Mediated by Wnt1 Induction in Osteoblast-Lineage Cells. (2022). 10.1101/2022.02.28.482178.

61. L. Y. Lawson, et al., Osteoblast-Specific Wnt Secretion Is Required for Skeletal Homeostasis and Loading-Induced Bone Formation in Adult Mice. J Bone Miner Res 37, 108–120 (2022).

62. T. Ganesh, L. E. Laughrey, M. Niroobakhsh, N. Lara-Castillo, Multiscale finite element modeling of mechanical strains and fluid flow in osteocyte lacunocanalicular system. Bone 137, 115328 (2020).

63. J. D. Currey, Stress concentrations in bone. Journal of Cell Science S3-103, 111–133 (1962).

64. R. P. Main, et al., Murine Axial Compression Tibial Loading Model to Study Bone Mechanobiology: Implementing the Model and Reporting Results. J Orthop Res 38, 233– 252 (2020).

65. A. I. Birkhold, et al., Mineralizing surface is the main target of mechanical stimulation independent of age: 3D dynamic in vivo morphometry. Bone 66, 15–25 (2014).

66. Q. A. Meslier, N. Migotsky, S. N. Lamia, E. L. Scheller, M. J. Silva, Protocol for 3D digital dynamic histomorphometry of mouse bone via time-lapse registration of serial microCT scans. STAR Protocols 7, 104787 (2026).

67. M. D. Brodt, M. J. Silva, Aged mice have enhanced endocortical response and normal periosteal response compared with young-adult mice following 1 week of axial tibial compression. J Bone Miner Res 25, 2006–2015 (2010).

68. M. M. Moran, et al., Intramembranous bone regeneration differs among common inbred mouse strains following marrow ablation. Journal Orthopaedic Research 33, 1374–1381 (2015).

69. D. W. Huang, B. T. Sherman, R. A. Lempicki, Bioinformatics enrichment tools: paths toward the comprehensive functional analysis of large gene lists. Nucleic Acids Research 37, 1–13 (2009).

70. D. W. Huang, B. T. Sherman, R. A. Lempicki, Systematic and integrative analysis of large gene lists using DAVID bioinformatics resources. Nat Protoc 4, 44–57 (2009).

71. P. Langfelder, S. Horvath, WGCNA: an R package for weighted correlation network analysis. BMC Bioinformatics 9, 559 (2008).

72. P. Langfelder, R. Luo, M. C. Oldham, S. Horvath, Is my network module preserved and reproducible? PLoS Comput Biol 7, e1001057 (2011).

