## Supplementary methods for "Bone Response to Mechanical Loading is Highly Heritable and Correlates with Osteocyte Lacunar Morphology in Genetically Diverse Mouse Strains"

### **Supplemental Methods**

#### Mouse housing and attrition

Inbred mice were group-housed, with up to 5 individuals per cage. Male DO mice were housed individually, while female DO mice were housed in groups of up to 3 individuals per cage, following recommendations from JAX to minimize intra-group aggression. Mice were allowed ad libitum access to chow and water. A total of 188 inbred and 50 DO mice were enrolled in the study. Of these, 19 mice died during the study period (2 PWK, 2 A/J, 2 WSB, 1 CAST, 2 129S1, 4 NZO, 5 NOD, and 1 DO). One WSB mouse escaped from its cage and was excluded from the study. Final analysis therefore included 169 inbred and 49 DO mice.

#### Dynamic histomorphometry

Briefly, tibias were cleaned of muscle and stored in 70% ethanol until plastic embedding. Bones were dehydrated in ethanol and embedded in methyl methacrylate (MMA, Thermo Scientific AAA130300F). Blocks were cut transversely (Leica SP1600 Microtome) at the site of peak strain, 5 mm proximal to the TFJ; three 100 µm thick slices were taken. Sections were imaged on a confocal microscope (Leica TCS SPEII, 10x lens, 2048x2048, 400Hz, 1.5x zoom) with a pixel resolution of 0.36 µm. A maximum projection was created from a 5 slice, 20 µm thick z-stack for final analysis. Images were blinded and fluorochrome-labeled surfaces were analyzed (Osteo II, Bioquant) separately on the periosteal and endocortical surfaces (1). In cases where no double label was visualized, a value of 0.3 µm/day was assigned for the respective mineral apposition rate (MAR) (2, 3). Relative values for all outcomes were calculated as loaded minus non-loaded.

#### RNA sequencing of bone tissue

RNA integrity (RIN) was determined using Agilent Bioanalyzer. Library preparation was performed with 100ng to 500ng of total RNA. Libraries were generated with Watchmaker Library Prep Kit with Polaris Depletion (Watchmaker). Only samples that passed library quality control (QC) metrics were sequenced. Sequencing was performed on an Illumina NovaSeq 6000 using paired end reads extending 150 bases. Basecalls and demultiplexing were performed with Illumina's bcl2fastq software and a custom python demultiplexing program with a maximum of one mismatch in the indexing read.  RNA-seq reads were then aligned to the Ensembl release

101 primary assembly with STAR version 2.7.9a (4). Gene counts were derived from the number of uniquely aligned unambiguous reads by Subread:featureCount version 2.0.3 (5). Sequencing performance was assessed for the total number of aligned reads, total number of uniquely aligned reads, and features detected. The ribosomal fraction, known junction saturation, and read distribution over known gene models were quantified with RSeQC version 4.0 (6).

We used DESeq2 in R/Bioconductor with Wald test to determine the differentially expressed genes (DEGs) between conditions (7). First, principal component analysis (PCA) was performed with the top 500 most variable genes across samples to visualize sample clustering. Based on differences in within-group variability, count data was subset by timepoint, and loaded samples were compared to nonloaded controls at each timepoint while controlling inter-animal variability. All count matrices were filtered to remove low-expression genes; only genes that have CPM ≥ 1 in at least one sample were included in the analysis. p-values were adjusted for multiple testing using the Benjamini and Hochberg method, and differential expression significance was set at p-adjusted (p-adj) $< 0.05$ and |linear fold change| $> 1.5$. RNAseq of 129S1 was compared to our published data set in age- and sex-matched B6 mice (8).

<u>Co-expression network analysis</u>

Weighted gene co-expression network analysis (WGCNA) was performed on 5-day 129S1 samples from the current study and the historical 5-day B6 samples (9). Briefly, TMM normalization of gene counts was performed in EdgeR in R/Bioconductor to adjust for differences in library size between samples (10). Normalized counts were then transformed to log2 counts-per-million using Limma voomWithQualityWeights (11), and the resulting expression matrix was input to WGCNA in R. Separate signed hybrid networks were constructed for 129S1 and B6 samples. All genes were correlated across each other by Pearson correlations and clustered by expression similarity into modules using a power threshold empirically determined from the data. An eigengene was then created for each de novo cluster and its expression profile was then correlated with loading (loaded vs nonloaded) within each network. Module preservation was tested to assess how well network structure of 129S1 modules were preserved in B6 dataset by determining Z summary and median rank (12). To identify modules of genes enriched for BMD GWAS genes, we obtained the mouse homolog list curated by (13) and used Fisher's exact test to measure the overrepresentation of GWAS genes in each module of 129S1 and B6 separately. We then applied Benjamini-Hochberg correction to correct for multiple testing with significance cutoff of 0.05.

**References**


1. M. D. Brodt, M. J. Silva, Aged mice have enhanced endocortical response and normal periosteal response compared with young-adult mice following 1 week of axial tibial compression. *J Bone Miner Res* **25**, 2006–2015 (2010).
2. D. W. Dempster, *et al.*, Standardized nomenclature, symbols, and units for bone histomorphometry: a 2012 update of the report of the ASBMR Histomorphometry Nomenclature Committee. *J Bone Miner Res* **28**, 2–17 (2013).
3. M. R. Allen, D. B. Burr, “Techniques in Histomorphometry” in *Basic and Applied Bone Biology*, (Elsevier, 2019), pp. 141–158.
4. A. Dobin, *et al.*, STAR: ultrafast universal RNA-seq aligner. *Bioinformatics* **29**, 15–21 (2013).
5. Y. Liao, G. K. Smyth, W. Shi, featureCounts: an efficient general purpose program for assigning sequence reads to genomic features. *Bioinformatics* **30**, 923–930 (2014).
6. L. Wang, S. Wang, W. Li, RSeQC: quality control of RNA-seq experiments. *Bioinformatics* **28**, 2184–2185 (2012).
7. M. I. Love, W. Huber, S. Anders, Moderated estimation of fold change and dispersion for RNA-seq data with DESeq2. *Genome Biology* **15**, 550 (2014).
8. C. J. Chermside-Scabbo, *et al.*, Old Mice Have Less Transcriptional Activation But Similar Periosteal Cell Proliferation Compared to Young-Adult Mice in Response to in vivo Mechanical Loading. *J Bone Miner Res* **35**, 1751–1764 (2020).
9. P. Langfelder, S. Horvath, WGCNA: an R package for weighted correlation network analysis. *BMC Bioinformatics* **9**, 559 (2008).
10. M. D. Robinson, D. J. McCarthy, G. K. Smyth, edgeR: a Bioconductor package for differential expression analysis of digital gene expression data. *Bioinformatics* **26**, 139–140 (2010).
11. R. Liu, *et al.*, Why weight? Modelling sample and observational level variability improves power in RNA-seq analyses. *Nucleic Acids Res* **43**, e97 (2015).

12. P. Langfelder, R. Luo, M. C. Oldham, S. Horvath, Is my network module preserved and reproducible? *PLoS Comput Biol* **7**, e1001057 (2011).

13. O. L. Sabik, G. M. Calabrese, E. Taleghani, C. L. Ackert-Bicknell, C. R. Farber, Identification of a Core Module for Bone Mineral Density through the Integration of a Co-expression Network and GWAS Data. *Cell Rep* **32**, 108145 (2020).
