## Supplementary figures and tables for "Bone Response to Mechanical Loading is Highly Heritable and Correlates with Osteocyte Lacunar Morphology in Genetically Diverse Mouse Strains"

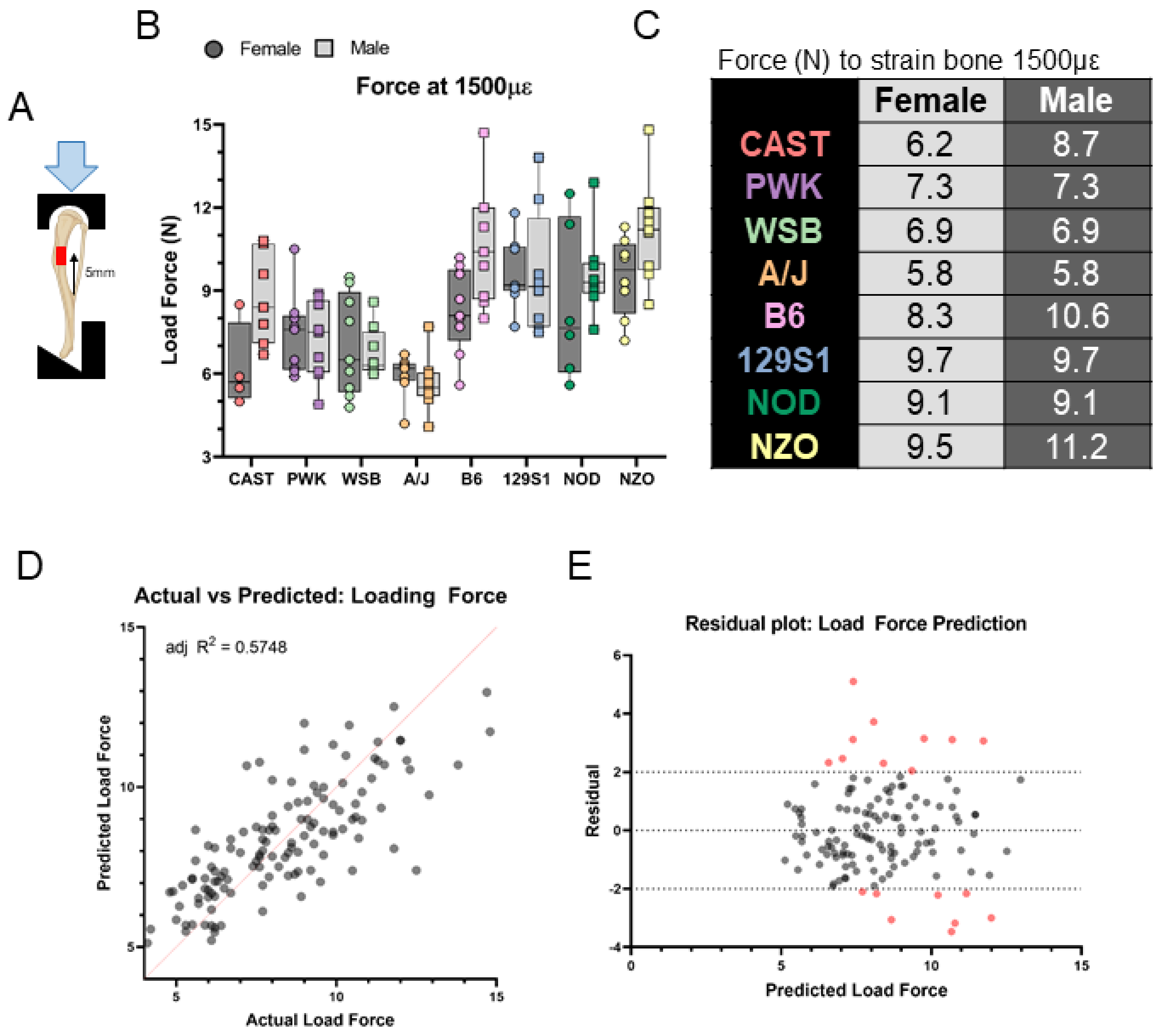


Force (N) to strain bone 1500με

| | Female | Male |
|---|---|---|
| CAST | 6.2 | 8.7 |
| PWK | 7.3 | 7.3 |
| WSB | 6.9 | 6.9 |
| A/J | 5.8 | 5.8 |
| B6 | 8.3 | 10.6 |
| 129S1 | 9.7 | 9.7 |
| NOD | 9.1 | 9.1 |
| NZO | 9.5 | 11.2 |

$$Force = 14.98 - 5.38(T.Ar) + 24.1(Ct.Ar) - 0.07(Weight) - 13.68(AreaRatio) - 10.98(Eccentricity$$

**Fig. S1: Mice were strain-gauged to determine equivalent load to produce 1500 μƐ.** (A) Strain gauge was glued 5 mm proximal to the distal tibiofibular junction on the anteromedial surface and strains recorded during tibial loading. (B) Force to engender 1500 με was calculated for non-loaded Inbred Founder strain mice. (C) The average force per strain/sex group that was used to load animals in this study. (D) Results from the multi-variable linear regression showing the predicted loading force from the model compared to the actual load force calculated from strain gauging. (E) Residuals are mainly between -2N and +2N.

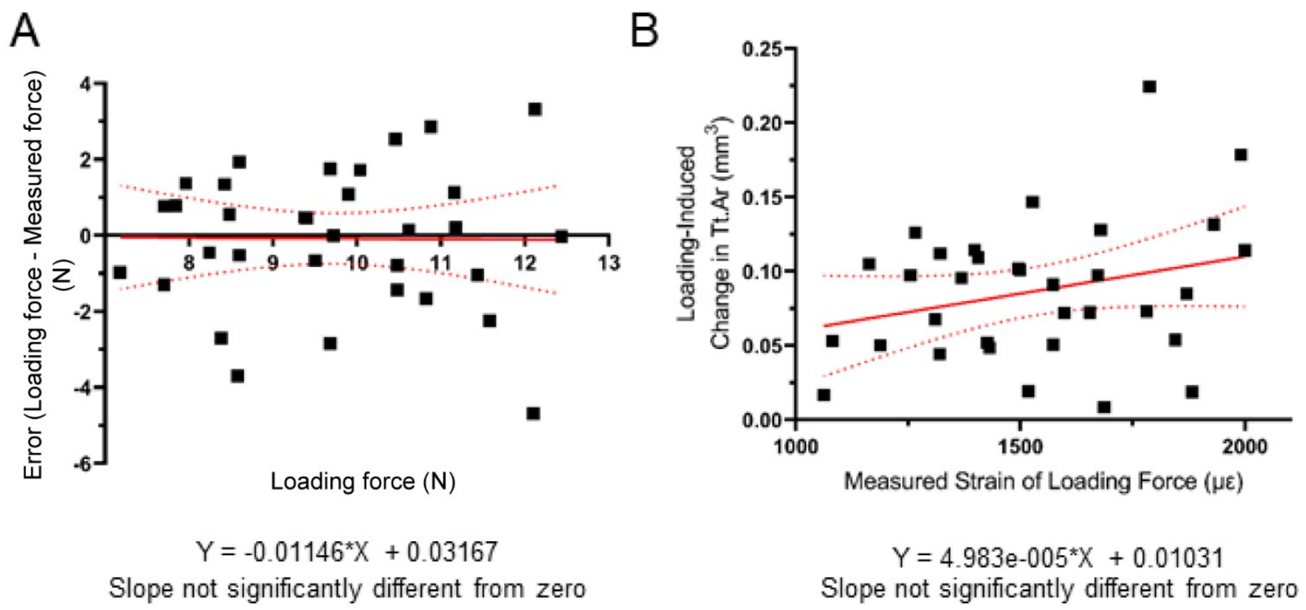


**Fig. S2: Loading force determination error in diversity outbred (DO) mice.** Loading forces for DO mice were calculated using the equation shown in Fig. S1 to engender 1500 μƐ and were applied to the right limb. To validate these calculated forces, strain gauging was performed postmortem on the contralateral, non-loaded tibia of DO mice to measure the force required to engender 1500 μƐ. (A) The errors in loading force are evenly distributed across the range of loading forces. The regression slope is not different from zero. (B) Correlation of the strain DO mice were loaded with loading-induced change in total area. The slope is not different from zero indicating there is no relationship between the error in loading force and bone formation.

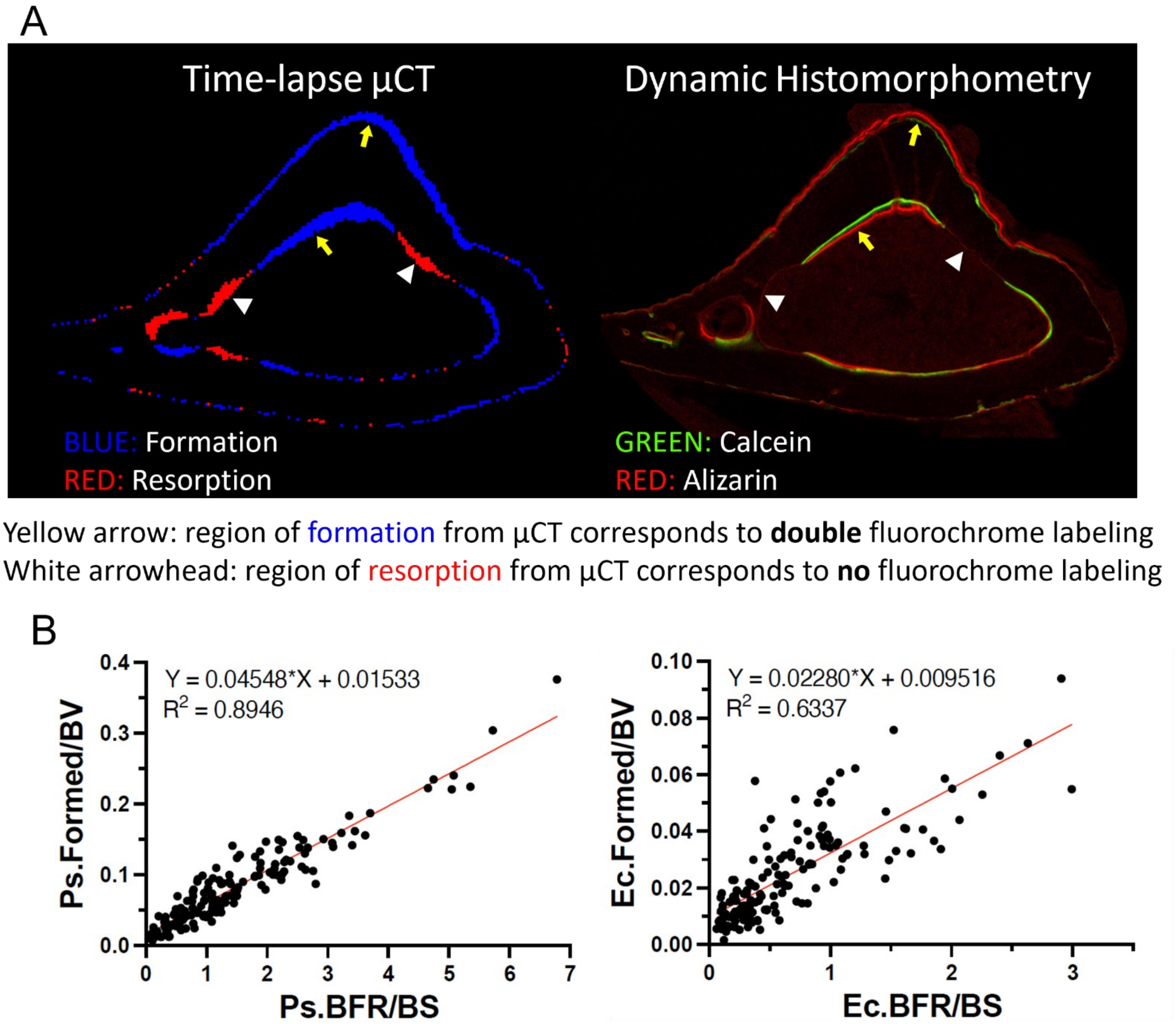


**Fig. S3: Validation of image registration.** (A) Left: Example of time-lapse microCT image showing formed (blue) and resorbed (red) bone voxels. Right: Fluorescent image of the same sample used for dynamic histomorphometry analysis. There is good qualitative agreement between the two images, i.e., site of thickest formed bone match the sites of where there is double fluorochrome labeled surface, while sites of resorbed bone match sites with no fluorochrome labeled surface. (B) Linear regression shows strong correlation between formed bone volume and bone formation rate on the periosteal surface, and moderate correlations on the endocortical surface. Modified from Meslier et al., https://doi.org/10.64898/2026.05.28.728585.

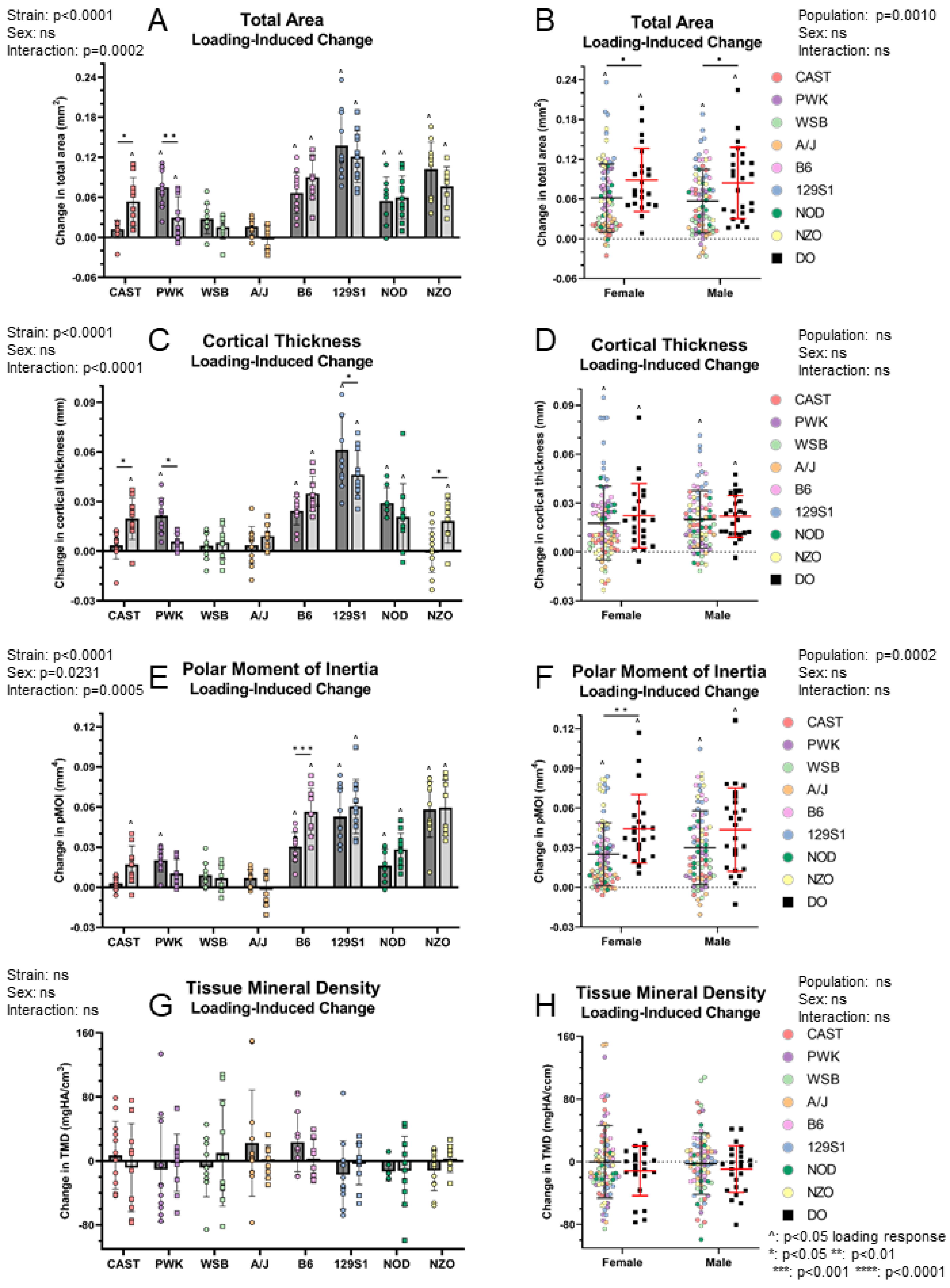
Strain: p<0.0001
Sex: ns
Interaction: p=0.0002
A
Total Area
Loading-Induced Change
Change in total area (mm$^2$)
0.24
0.18
0.12
0.06
0.00
-0.06
CAST
PWK
WSB
A/J
B6
129S1
NOD
NZO
B
Total Area
Loading-Induced Change
Population: p=0.0010
Sex: ns
Interaction: ns
Female
Male
CAST
PWK
WSB
A/J
B6
129S1
NOD
NZO
DO
Strain: p<0.0001
Sex: ns
Interaction: p<0.0001
C
Cortical Thickness
Loading-Induced Change
Change in cortical thickness (mm)
0.09
0.06
0.03
0.00
-0.03
D
Cortical Thickness
Loading-Induced Change
Population: ns
Sex: ns
Interaction: ns
Strain: p<0.0001
Sex: p=0.0231
Interaction: p=0.0005
E
Polar Moment of Inertia
Loading-Induced Change
Change in pMOI (mm$^4$)
0.12
0.09
0.06
0.03
0.00
-0.03
F
Polar Moment of Inertia
Loading-Induced Change
Population: p=0.0002
Sex: ns
Interaction: ns
Strain: ns
Sex: ns
Interaction: ns
G
Tissue Mineral Density
Loading-Induced Change
Change in TMD (mgHA/cm$^3$)
160
80
0
-80
H
Tissue Mineral Density
Loading-Induced Change
Population: ns
Sex: ns
Interaction: ns
Change in TMD (mgHA/ccm)
^: p<0.05 loading response
*: p<0.05 **: p<0.01
***: p<0.001 ****: p<0.0001

**Fig. S4: Loading-induced changes in bone morphology vary with mouse strain.** Tibiae were unilaterally loaded daily for 2 wks, and both nonloaded and loaded tibiae were microCT-scanned at the beginning (baseline) and end of the study period (Fig 2A). Relative changes with time were calculated for each limb with respect to baseline (Δ), and then relative changes due to loading were calculated as differences between limbs (ΔΔ). (A) Relative changes in total area of eight inbred strains and (B) inbred strains (pooled) and diversity outbred (DO) mice. (C) Relative changes in cortical thickness of the inbred strains and (D) inbred strains (pooled) and DO mice. (E) Relative changes in polar moment of inertia (pMOI) of the inbred strains and (F) inbred strains (pooled) and DO mice. (G) Relative changes in tissue mineral density (TMD) of the inbred strains and (H) inbred strains (pooled) and DO mice. Two-way ANOVA (sex and strain or population). Data shown as mean±SD.

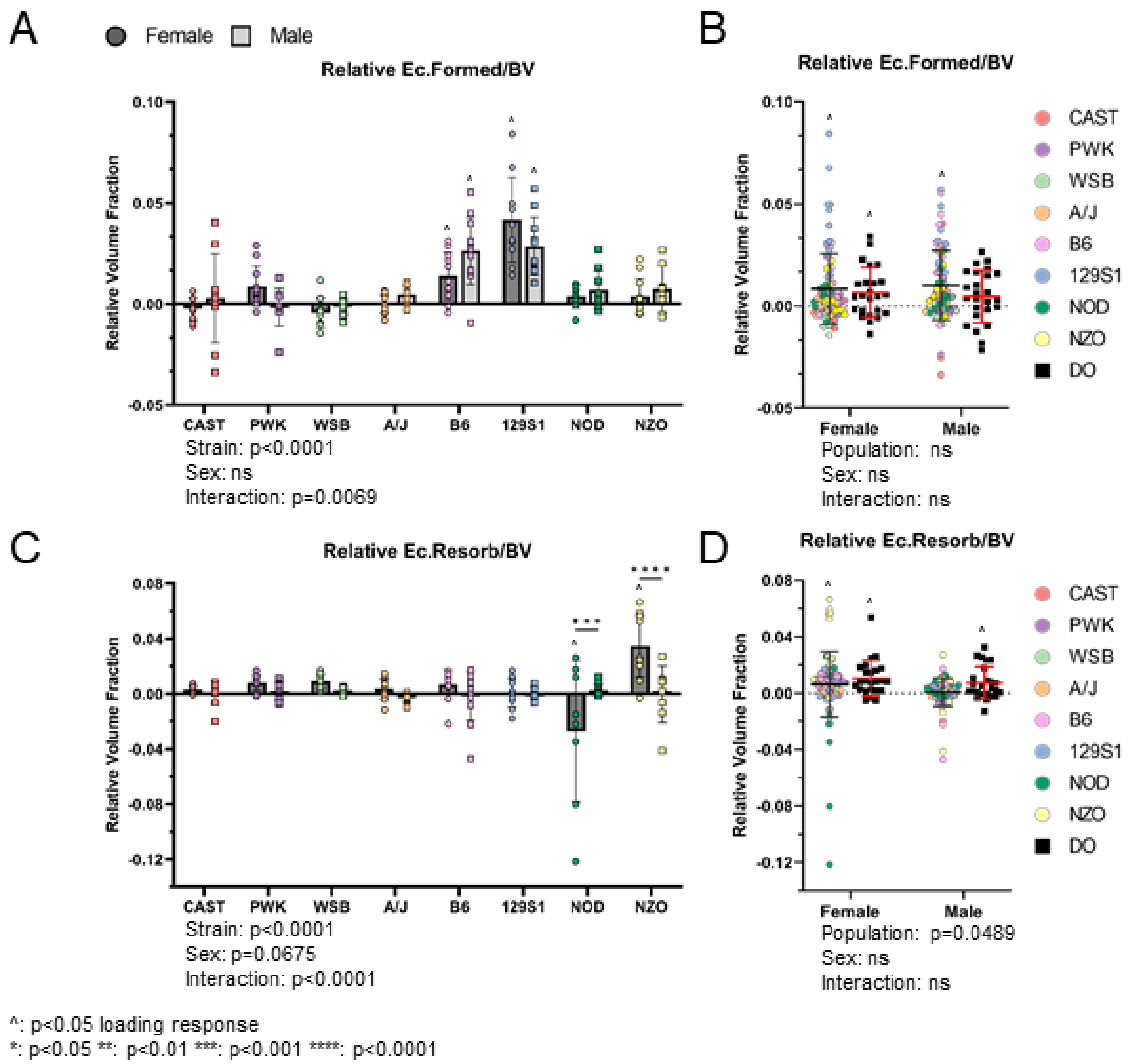


**Fig. S5: Loading-induced increases in endocortical formed bone volume vary with mouse strain.** The pre- and post-loading tibia microCT scans were registered in 3D to obtain formed and resorbed bone volumes. Relative changes were calculated with respect to nonloaded tibia. (A) Relative endocortical (Ec.) formed volume fraction in eight inbred strains and (B) in inbred strains (pooled) and diversity outbred (DO) mice. (C) Relative endocortical resorbed volume fraction in the inbred strains and (D) in inbred strains (pooled) and DO mice. Two-way ANOVA (sex and strain or population). Data shown as mean±SD.

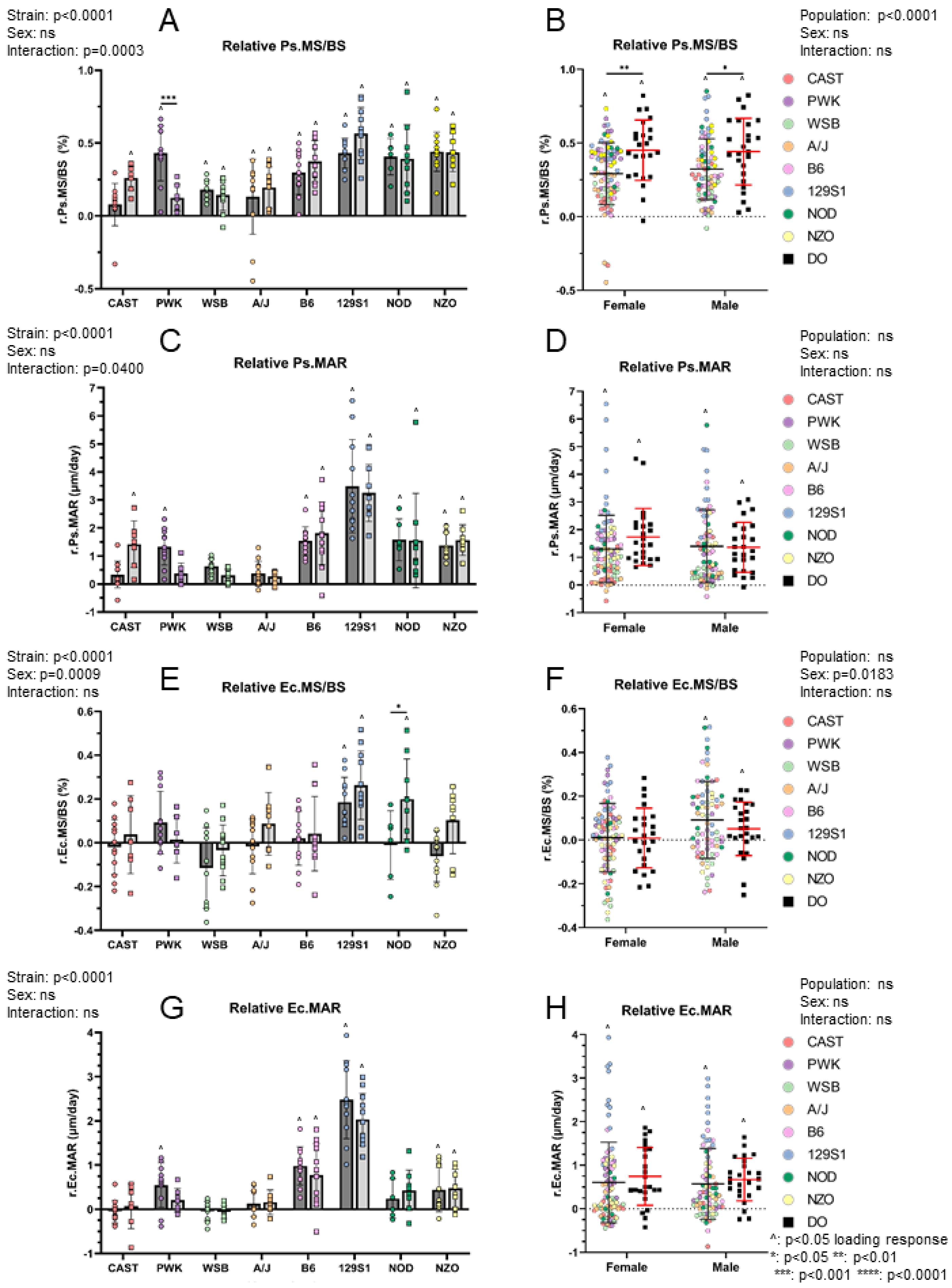
Strain: p<0.0001
Sex: ns
Interaction: p=0.0003
A
Relative Ps.MS/BS
r.Ps.MS/BS (%)
CAST
PWK
WSB
A/J
B6
129S1
NOD
NZO
B
Relative Ps.MS/BS
Population: p<0.0001
Sex: ns
Interaction: ns
r.Ps.MS/BS (%)
Female
Male
CAST
PWK
WSB
A/J
B6
129S1
NOD
NZO
DO
Strain: p<0.0001
Sex: ns
Interaction: p=0.0400
C
Relative Ps.MAR
r.Ps.MAR (μm/day)
D
Relative Ps.MAR
Population: ns
Sex: ns
Interaction: ns
r.Ps.MAR (μm/day)
Strain: p<0.0001
Sex: p=0.0009
Interaction: ns
E
Relative Ec.MS/BS
r.Ec.MS/BS (%)
F
Relative Ec.MS/BS
Population: ns
Sex: p=0.0183
Interaction: ns
r.Ec.MS/BS (%)
Strain: p<0.0001
Sex: ns
Interaction: ns
G
Relative Ec.MAR
r.Ec.MAR (μm/day)
H
Relative Ec.MAR
Population: ns
Sex: ns
Interaction: ns
r.Ec.MAR (μm/day)
^: p<0.05 loading response
*: p<0.05 **: p<0.01
***: p<0.001 ****: p<0.0001

**Fig. S6: Loading-induced increases in bone formation rate vary with mouse strain.** Bone formation indices were measured on nonloaded and loaded tibia using classical dynamic histomorphometry. Relative changes were calculated with respect to nonloaded tibia. (A) Relative periosteal (Ps.) mineralizing surface (MS/BS) in eight inbred strains and (B) in inbred strains (pooled) and diversity outbred (DO) mice. (C) Relative periosteal mineral apposition rate (MAR) in the inbred strains and (D) in inbred strains (pooled) and DO mice. (E) Relative endocortical (Ec.) mineralizing surface in inbred strains and (B) in inbred strains (pooled) and diversity outbred (DO) mice. (C) Relative endocortical mineral apposition rate in inbred strains and (D) in inbred strains (pooled) and DO mice. Two-way ANOVA (sex and strain or population). Data shown as mean±SD.

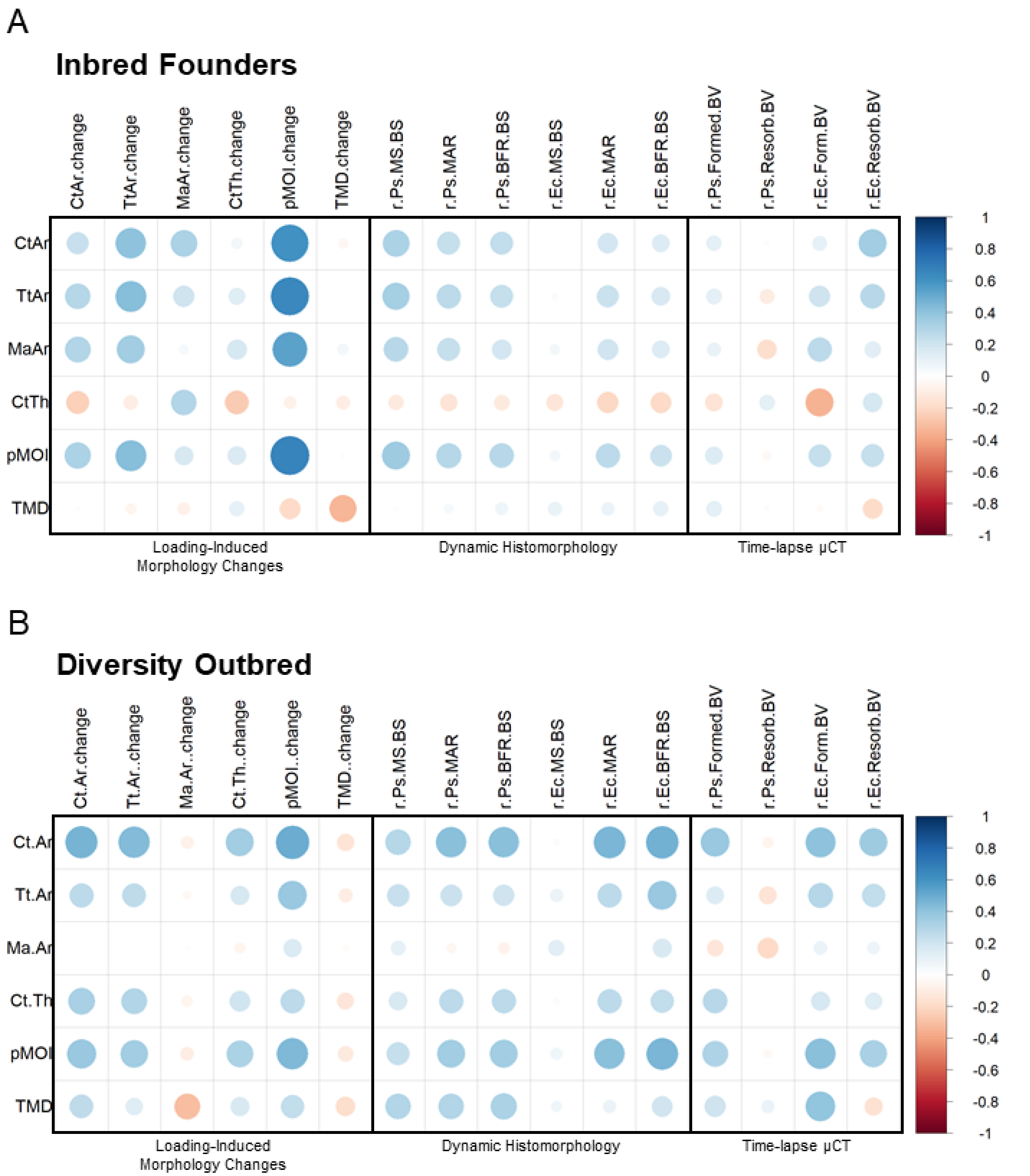


**Fig. S7: Correlations between starting μCT parameters (pre-loading) and loading response outcomes were not strong.** Pearson's correlation matrix between starting μCT parameters (pre-loading) and all loading response outcomes. Correlations in (A) inbred founder strains (n = 171) and (B) diversity outbred (DO) mice (n = 49).

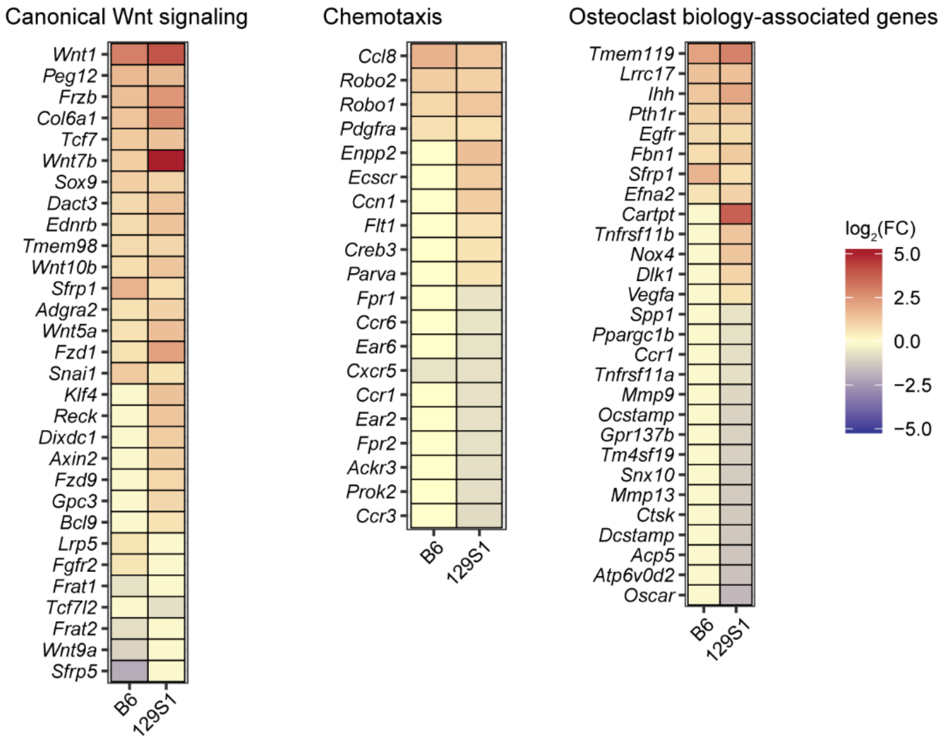


**Fig. S8: Expression levels of genes associated with specific gene ontology biological processes.**

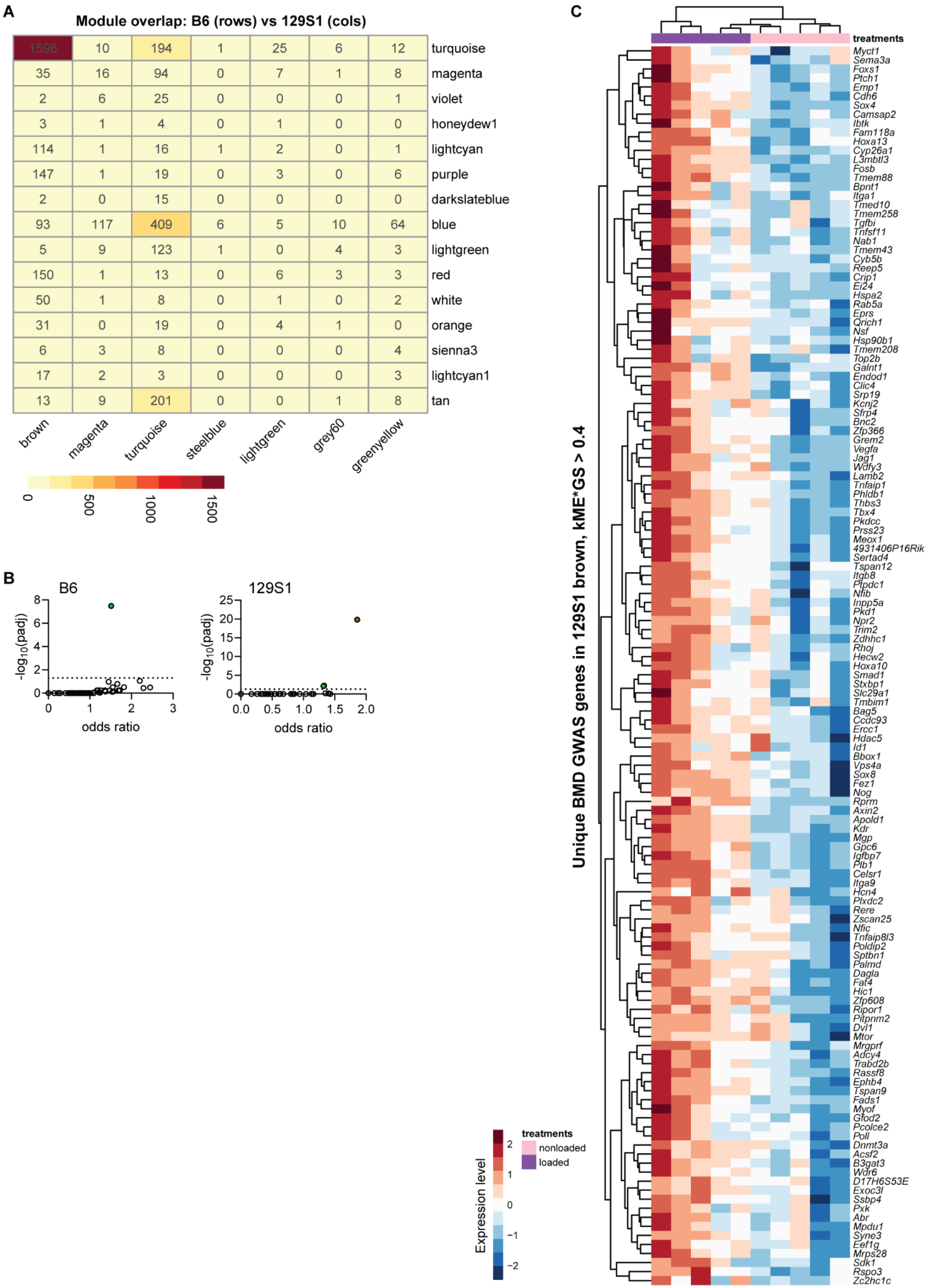
A
Module overlap: B6 (rows) vs 129S1 (cols)
1598 10 194 1 25 6 12 turquoise
35 16 94 0 7 1 8 magenta
2 6 25 0 0 0 1 violet
3 1 4 0 1 0 0 honeydew1
114 1 16 1 2 0 1 lightcyan
147 1 19 0 3 0 6 purple
2 0 15 0 0 0 0 darkslateblue
93 117 409 6 5 10 64 blue
5 9 123 1 0 4 3 lightgreen
150 1 13 0 6 3 3 red
50 1 8 0 1 0 2 white
31 0 19 0 4 1 0 orange
6 3 8 0 0 0 4 sienna3
17 2 3 0 0 0 3 lightcyan1
13 9 201 0 0 1 8 tan
brown
magenta
turquoise
steelblue
lightgreen
grey60
greenyellow
0
500
1000
1500
B
B6
-log10(padj)
odds ratio
129S1
C
treatments
Unique BMD GWAS genes in 129S1 brown, kME*GS > 0.4
Myct1
Sema3a
Foxs1
Ptch1
Emp1
Cdh6
Sox4
Camsap2
Ibtk
Fam118a
Hoxa13
Cyp26a1
L3mbtl3
Fosb
Tmem88
Bpnt1
Itga1
Tmed10
Tmem258
Tgfbi
Tnfsf11
Nab1
Tmem43
Cyb5b
Reep5
Crip1
Ei24
Hspa2
Rab5a
Eprs
Qrich1
Nsf
Hsp90b1
Tmem208
Top2b
Galnt1
Endod1
Clic4
Srp19
Kcnj2
Sfrp4
Bnc2
Zfp366
Grem2
Vegfa
Jag1
Wdfy3
Lamb2
Tnfaip1
Phldb1
Thbs3
Tbx4
Pkdcc
Prss23
Meox1
4931406P16Rik
Sertad4
Tspan12
Itgb8
Ptpdc1
Nfib
Inpp5a
Pkd1
Npr2
Trim2
Zdhhc1
Rhoj
Hecw2
Hoxa10
Smad1
Stxbp1
Slc29a1
Tmbim1
Bag5
Ccdc93
Ercc1
Hdac5
Id1
Bbox1
Vps4a
Sox8
Fez1
Nog
Rprm
Axin2
Apold1
Kdr
Mgp
Gpc6
Igfbp7
Plb1
Celsr1
Itga9
Hcn4
Plxdc2
Rere
Zscan25
Nfic
Tnfaip8l3
Poldip2
Sptbn1
Palmd
Dagla
Fat4
Hic1
Zfp608
Ripor1
Pitpnm2
Dvl1
Mtor
Mrgprf
Adcy4
Trabd2b
Rassf8
Ephb4
Tspan9
Fads1
Myof
Gfod2
Pcolce2
Poll
Dnmt3a
Acsf2
B3gat3
Wdr6
D17H6S53E
Exoc3l
Ssbp4
Pxk
Abr
Mpdu1
Syne3
Eef1g
Mrps28
Sdk1
Rspo3
Zc2hc1c
Expression level
treatments
nonloaded
loaded

**Fig. S9: Loading response is largely conserved between B6 and 129S1 at transcriptome level.** (A) Gene overlap matrix between significant ($p<0.1$) B6 and 129S1 modules. (B) Module enrichments for genes overlapping a human BMD GWAS association (Fisher's exact test). (C) 129S1-unique GWAS genes.

**Table S1: Select gene ontology biological processes enriched among unique and shared DEGs after 5-day loading in B6 and 129S1. FDR<0.2.**

**B6 unique genes**

| Term | Count | Pop.Hits | Fold.Enrichment |
|---|---|---|---|
| gluconeogenesis | 14 | 62 | 9.77 |
| extracellular matrix organization | 17 | 157 | 4.69 |
| response to cold | 9 | 37 | 10.53 |
| lipid metabolic process | 42 | 800 | 2.27 |
| transmembrane transport | 33 | 561 | 2.55 |
| chondrocyte differentiation | 10 | 71 | 6.09 |
| bone development | 11 | 97 | 4.91 |
| angiogenesis | 19 | 302 | 2.72 |
| positive regulation of transcription by RNA polymerase II | 52 | 1328 | 1.69 |
| positive regulation of JNK cascade | 10 | 115 | 3.76 |
| cell-cell adhesion | 14 | 220 | 2.75 |
| cellular response to transforming growth factor beta stimulus | 8 | 78 | 4.44 |
| cellular response to hypoxia | 11 | 149 | 3.19 |
| peripheral nervous system myelin formation | 3 | 4 | 32.45 |
| regulation of cell population proliferation | 10 | 210 | 2.06 |

**129S1 unique genes**

| Term | Count | Pop.Hits | Fold.Enrichment |
|---|---|---|---|
| extracellular matrix organization | 30 | 157 | 6.93 |
| angiogenesis | 33 | 302 | 3.97 |
| cell-cell adhesion | 26 | 220 | 4.29 |
| cell migration | 28 | 328 | 3.1 |
| monoatomic ion transport | 40 | 635 | 2.28 |
| skeletal system development | 16 | 135 | 4.3 |
| negative regulation of canonical Wnt signaling pathway | 16 | 140 | 4.14 |
| bone mineralization | 11 | 67 | 5.96 |
| chemotaxis | 15 | 143 | 3.8 |
| positive regulation of BMP signaling pathway | 8 | 40 | 7.26 |
| osteoblast differentiation | 13 | 119 | 3.96 |
| nervous system development | 30 | 496 | 2.2 |
| positive regulation of ERK1 and ERK2 cascade | 17 | 222 | 2.78 |
| positive regulation of epithelial cell proliferation | 10 | 91 | 3.98 |
| smoothened signaling pathway | 11 | 112 | 3.56 |
| neuropeptide signaling pathway | 11 | 112 | 3.56 |
| Wnt signaling pathway | 17 | 244 | 2.53 |
| positive regulation of transcription by RNA polymerase II | 56 | 1328 | 1.53 |
| positive regulation of apoptotic process | 22 | 388 | 2.06 |
| osteoclast differentiation | 7 | 55 | 4.62 |

**Shared genes**

| Term | Count | Pop.Hits | Fold.Enrichment |
|---|---|---|---|
| collagen fibril organization | 31 | 63 | 20.3 |
| extracellular matrix organization | 27 | 157 | 7.09 |
| osteoblast differentiation | 22 | 119 | 7.63 |
| angiogenesis | 31 | 302 | 4.23 |
| bone development | 17 | 97 | 7.23 |
| cell migration | 29 | 328 | 3.65 |
| nervous system development | 35 | 496 | 2.91 |
| bone mineralization | 13 | 67 | 8 |
| canonical Wnt signaling pathway | 16 | 113 | 5.84 |
| cell-cell adhesion | 21 | 220 | 3.94 |
| chondrocyte differentiation | 12 | 71 | 6.97 |
| monoatomic ion transport | 36 | 635 | 2.34 |
| negative regulation of canonical Wnt signaling pathway | 15 | 140 | 4.42 |
| positive regulation of mesenchymal cell proliferation | 8 | 36 | 9.17 |
| ephrin receptor signaling pathway | 8 | 43 | 7.68 |
| negative chemotaxis | 7 | 42 | 6.88 |
| Wnt signaling pathway, planar cell polarity pathway | 5 | 20 | 10.31 |
| potassium ion transport | 12 | 153 | 3.24 |
| positive regulation of gene expression | 28 | 593 | 1.95 |
| sodium ion transport | 12 | 155 | 3.19 |
| integrin-mediated signaling pathway | 10 | 111 | 3.72 |
| fat cell differentiation | 10 | 112 | 3.68 |
| smoothened signaling pathway | 10 | 112 | 3.68 |
| calcium ion transmembrane transport | 11 | 148 | 3.07 |
| semaphorin-plexin signaling pathway | 6 | 43 | 5.76 |
| BMP signaling pathway | 9 | 104 | 3.57 |

**Table S2: Top 5 GO biological processes enriched in loading associated modules.**

**B6**

| Module | Gene ontology biological process | p-adj | fold enrichment |
|---|---|---|---|
| Turquoise (+) | collagen fibril organization | 4.8E-22 | 6.88 |
| | nervous system development | 1.3E-20 | 2.64 |
| | vesicle-mediated transport | 1.7E-17 | 2.94 |
| | cell differentiation | 1.7E-17 | 1.96 |
| | protein transport | 2.0E-17 | 2.25 |
| Magenta (-) | myelination | 8.3E-05 | 8.69 |
| | positive regulation of gene expression | 3.0E-04 | 3.14 |
| | actin cytoskeleton organization | 7.9E-03 | 4.12 |
| | lipid metabolic process | 1.3E-02 | 2.41 |
| | response to insulin | 1.3E-02 | 6.42 |
| Violet (-) | *no significant enrichment* | | |

**129S1**

| Module | Gene ontology biological process | p-adj | fold enrichment |
|---|---|---|---|
| Brown (+) | angiogenesis | 7.0E-32 | 3.51 |
| | cell adhesion | 3.4E-28 | 2.49 |
| | extracellular matrix organization | 3.7E-24 | 4.15 |
| | collagen fibril organization | 3.3E-21 | 6.03 |
| | vesicle-mediated transport | 2.2E-20 | 2.89 |
| Magenta (-) | *no significant enrichment* | | |
| Turquoise (-) | regulation of transcription by RNA polymerase II | 1.5E-30 | 2.12 |
| | regulation of DNA-templated transcription | 8.1E-18 | 1.84 |
| | methylation | 1.2E-07 | 2.93 |
| | chromatin organization | 2.0E-03 | 1.83 |
| | chromatin remodeling | 5.2E-03 | 1.95 |

**Table S3: Enrichment of mouse homologs of human BMD GWAS.**

| B6 modules | module_size | gwas_genes_in_module | gwas_genes_in_network |
|---|---|---|---|
| turquoise | 2885 | 348 | 1501 |
| magenta | 457 | 48 | 1501 |
| purple | 420 | 38 | 1501 |
| lightcyan | 284 | 20 | 1501 |
| violet | 93 | 10 | 1501 |
| darkslateblue | 44 | 4 | 1501 |
| honeydew1 | 30 | 0 | 1501 |

| 129S1 modules | module_size | gwas_genes_in_module | gwas_genes_in_network |
|---|---|---|---|
| brown | 3550 | 391 | 1520 |
| magenta | 356 | 21 | 1520 |
| steelblue | 42 | 1 | 1520 |
| lightgreen | 149 | 6 | 1520 |
| turquoise | 4448 | 191 | 1520 |
